# IOP-independent treatment for neurodegeneration in glaucoma using optimized ROS-scavenging microparticles for sustained release of erythropoietin

**DOI:** 10.64898/2026.10.29.762812

**Authors:** Carlisle R. DeJulius, Sarah Naguib, Patricia Poley, Marselle Rasdall, Thomas Dean, Amy Connor, Sariah D’Empaire-Salomon, Rachel Hardin, Elisabeth Artis, Richard d’Arcy, Brian Carlson, David J. Calkins, Craig L. Duvall, Tonia S. Rex

## Abstract

Glaucoma is a slow-progressing neurodegenerative disease of the optic projection to the brain and a leading cause of blindness worldwide. Current treatments are targeted towards lowering intraocular pressure (IOP), the only known, modifiable risk factor; however, this strategy is ineffective in many patients. An early hallmark of glaucomatous neurodegeneration is increased reactive oxygen species (ROS), representing an alternative opportunity for intervention to halt or slow disease progression. In this work, we optimize a ROS-scavenging copolymer, poly(propylene sulfide-co-ethylene sulfide) (PPSES), for formulation of size-controlled, stabilized, antioxidant microparticles (MPs) for sustained release of the protein erythropoietin (EPO). EPO promotes biological antioxidant defense via activation of NRF2-mediated transcription of antioxidant genes. This MP system overcomes the short half-life of EPO and leverages the combined benefits of bioactive EPO with long-lasting, inherently antioxidant PPSES MPs. The lead EPO-MP formulation preserves neurons in both mouse and non-human primate models of glaucoma through suppression of oxidative stress and optic nerve degeneration. EPO-loaded PPSES MPs represent a translationally relevant therapy for long-term management of glaucoma independent of IOP.

## INTRODUCTION

Glaucoma is a leading cause of blindness worldwide, with treatments largely focused on lowering intraocular pressure (IOP), either by pharmacological or surgical means (*1*). However, many patients continue to lose vision despite IOP management. Thus, glaucoma is a prime candidate for innovation of new therapies that provide long-term protection of the neural retina and optic nerve, the tissues ultimately irreversibly damaged in glaucoma. A hallmark of glaucoma pathogenesis is increased oxidative stress, both in the trabecular meshwork, affecting fluid outflow (*2*), and in the retina, reducing survival of retinal ganglion cells (RGCs) and their axons, which comprise the optic projection to the brain (*3–5*). Glaucomatous animal models also exhibit elevated reactive oxygen species (ROS) levels and reduced quantities of antioxidant proteins (*3*, *6*). The retina initiates an endogenous protective response to elevated IOP driven by activation of NRF2 and the antioxidant response element (ARE) (*7*). Further, inhibition of this pathway accelerates glaucoma pathogenesis, while supplementation with exogenous antioxidants can reduce glaucoma pathogenesis in animal models (*8*, *9*). Mechanistically, oxidative stress impacts glaucoma by causing a combination of axonopathy, increased IOP, and optic nerve degeneration, resulting in vision loss.

In this study, we sought to innovate a sustained release microparticle (MP) formulation with antioxidant action to directly address the increased ROS detected in the retina during early-stage glaucoma. The gold standard polymer class for MP drug delivery is the hydrolytically degradable polyesters, particularly poly(lactic-co-glycolic acid) (PLGA), which is the basis of over 20 approved clinical products (*10*, *11*). PLGA breaks down into the naturally-present metabolic byproducts lactic and glycolic acid; however, accumulation of acidic degradation byproducts can lower local pH (*12*), elicit foreign-body response (*13*), and activate immune cells (*14*). As an alternative, we previously developed the ROS-responsive polythioether poly(propylene sulfide) (PPS) for MP formulations, which exhibits a hydrophobic-to-hydrophilic switch upon reaction with ROS like hydrogen peroxide (H_2_O_2_) (*15*). Through this mechanism, PPS can locally sponge ROS, triggering cargo release and breakdown of PPS-based nano- and microparticle formulations (*16–19*). We have previously shown the ability of PPS MPs to reduce tissue damage in models of peripheral arterial disease (*20*), osteoarthritis (*21*), and traumatic optic neuropathy (*22*). However, a major limitation of PPS MPs is its amorphous character (*23*). Greater crystallinity is required for forming stable, 10-100 µm diameter MPs, which is necessary to avoid particle loss due to rapid coalescence, structural breakdown, or cellular phagocytosis. To solve this challenge, we synthesized a series of poly(propylene sulfide-co-ethylene sulfide) (PPSES) random copolymers incorporating propylene sulfide and ethylene sulfide (ES), a monomer that yields more crystalline polymers (*24*).

We chose to optimize PPSES MPs for sustained release of erythropoietin (EPO), a hormone and cytokine with neuroprotective activity (*25–28*). Sustained delivery of EPO from PLGA MPs has demonstrated improved functional recovery and tissue preservation in rodent models of peripheral nerve injury (*29*, *30*), reduced RGC apoptosis in a rat model of optic nerve crush (*31*), and protected the RGCs in a mouse model of glaucoma (*32*). Our previous work with PPS-EPO MPs in a mouse model of traumatic optic neuropathy showed that sustained EPO delivery preserved retina and optic nerve function while reducing ROS and inflammatory cytokines (*22*). Mechanistically, we previously demonstrated that EPO activates NRF2 in a mouse model of glaucoma, resulting in increased production of antioxidant enzymes including superoxide dismutase 2 (SOD2), protection of the RGCs, and preservation of visual function (*32*). In this study, we optimized PPSES-EPO MPs toward a translatable product form and sought to test the system in a clinically-relevant non-human primate (NHP) model of glaucoma. Precise tuning of polymer composition and EPO formulation parameters yielded robust neuroprotection in both murine and nonhuman primate disease models.

## RESULTS

### Formulation of ROS-responsive MPs for sustained drug delivery

PPSES copolymers were synthesized according to **Scheme 1**. ES content ranged from 0 (PPS) to 50 mol% (**Supplementary Figure 1A-F**), above which the polymer crystallized during polymerization and was irrecoverable. Each polymer in the series was analyzed by GPC and found to have a molecular weight of approximately 15 kDa and polydispersity index (M_w_/M_n_) of 1.03-1.14 (**Supplementary Figure 2A-B**). Copolymerization with the ES monomer increased crystallinity of the polymers, with compositions containing >30% ES exhibiting broad exothermic crystallization peaks by differential scanning calorimetry (DSC) upon cooling from the melt (**Figure 1A**) and endothermic shifts under heating corresponding to melting of crystalline domains (**Supplementary Figure 2C**). PS_50_-co-ES_50_ exhibited the most robust crystallization, with T_c,onset_ of 47.5°. Successive self-nucleation (SSA) demonstrated temperature-dependent melting of crystallites in the PS_50_-co-ES_50_ polymer attributable to variability in the length of ES segments within the chain (Supplementary **Figure 2D**) (*23*). Further analysis via X-ray diffractive spectroscopy (XRD) detected a main peak at ∼20° in all compositions, typical ofamorphous polymers (*33*), and a secondary peak at ∼24° in PS_60_-co-ES_40_ and PS_50_-co-ES_50_, corresponding with the unit cell of ES (**Figure 1B**) (*24*). The percent crystallinity (X_c,%_) of PS_50_-co-ES_50_ and PS_60-_co-ES_40_ were calculated to be approximately 13 and 7%, respectively, from the area under the XRD crystallization peak.

**Figure 1:**
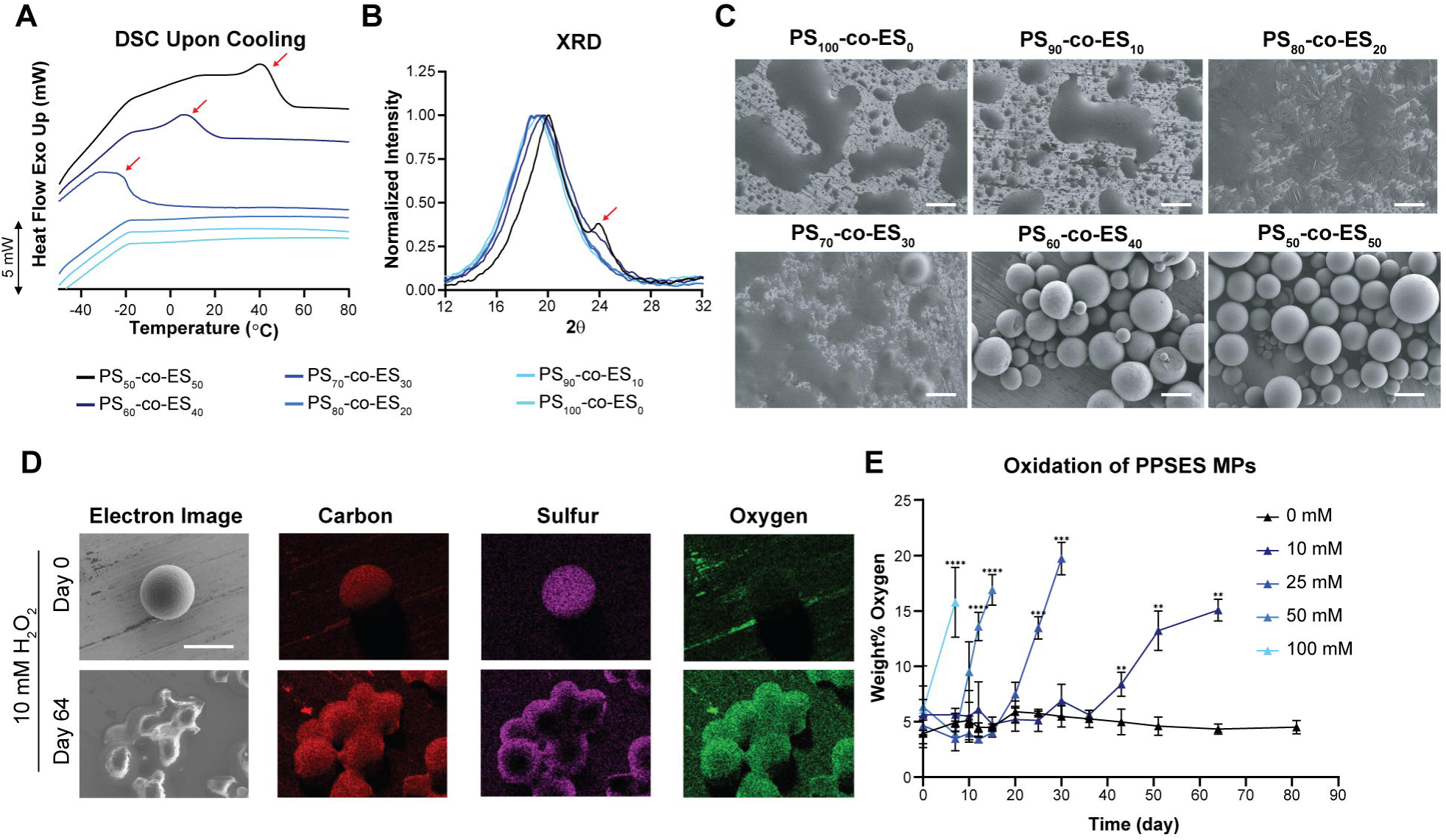
Characterization of PS-co-ES copolymer series. **(A)** Differential scanning calorimetry (DSC) showing exothermic (Exo) heat flow in the material during a temperature ramp, allowing identification of the crystallization process (red arrows). (B) X-ray diffractive spectroscopy (XRD) showing a crystalline peak at 20= 24° for the higher ES% (red arrow). **(C)** Scanning electron micrograph of PPSES microparticle (MP) series. Scale bar = 100 pm. **(D)** Elemental mapping via energy dispersive x-ray spectroscopy (EDS) of PS_50_-co-ES_50_ (PPSES) MPs exposed to 10 mM H_2_O_2_ demonstrates localization of the major constitutive elements of the MPs and allows visualization of polymer oxidation over time under exposure to H_2_O,. Red = carbon, purple = sulfur, and green = oxygen. Scale bar = 100 pm. (E) Quantification of oxygen weight % in the MPs over time after exposure to a range of concentrations of H_n_O_2_. Data shown for MPs >50 pm in diameter. **p<0.005, ***p<0.0005, ****p<0.0001.

**Scheme 1:**
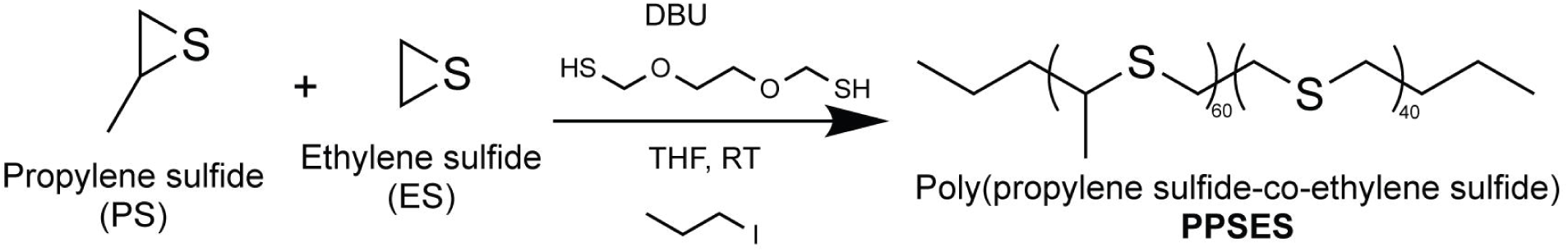
Synthesis of PPSES polymer series.

Formulation of MPs from the PPSES polymer series revealed that MP formation and stability were affected by PPSES copolymer composition. We developed an oil-in-water (O/W) emulsion protocol for formation of MPs with approximately 50-100 µm diameter. MP morphology was visualized by SEM, and ability of the MPs to be resuspended after lyophilization was also qualitatively assessed. Only PS_60_-co-ES_40_ and PS_50_-co-ES_50_ yielded stable MPs in this size range, while the remaining polymers had a coalesced appearance (**Figure 1C**). Qualitatively, the polymers with ES content less than 40 mol% formed viscous aggregates with poor handling characteristics relative to the more crystalline MP suspensions. The remainder of this work focuses on PS_50_-co-ES_50_, referred to henceforth as “PPSES”.

We utilized energy dispersive x-ray spectroscopy (EDS) to determine chemical oxidation kinetics of PPSES MPs (**Figure 1D-E** and **Supplementary Figure 3 and 4**). MPs are stable in aqueous conditions without ROS, with the morphology and oxygen percentage virtually unchanged even after 81 days of incubation (**Supplementary Figure 3 and 4** and **Figure 1E**). However, exposure to H_2_O_2_ causes gradual oxidation of the polymer, detectable by increased oxygen percentage; this process increases the water solubility of the polymers, resulting in eventual MP physical breakdown. In milder conditions (10 mM H_2_O_2_), there was a significant increase in oxygen percentage before gross morphology changes (**Figure 1E**). In all conditions, maximum oxidation occurred at the final time point before complete degradation (∼15-20% oxygen), and there was no difference in oxidation kinetics between “large” (>50 µm diameter, **Figure 1E**) and “small” (<50 µm diameter, **Supplementary Figure 4F**) MPs. Finally, PPSES MPs exhibited no cytotoxicity in ARPE-19 cells (**Supplementary Figure 2E**), as expected based on previous work with PPS (*22*), indicating their suitability for *in vivo* testing.

### Optimization of ROS-scavenging, EPO-loaded MPs in a mouse model of glaucoma

We formulated PPSES-EPO MPs using a standard water-in-oil-in-water (W/O/W) double emulsion process and a more exploratory solid-in-oil-in-water (S/O/W) double emulsion process. For the S/O/W formulation, EPO was first loaded into dextran nanoparticles (**Supplementary Figure 5A**) to protect the protein from interfacial degradation which occurs in W/O/W procedures (*34*). As EPO is a negatively charged protein (*35*), we incorporated positively charged diethylaminoethyl-dextran (DEAE-dextran), which increased encapsulation efficiency in the precursor dextran nanoparticles from 15.4% to 96.8% compared to dextran alone (**Supplementary Figure 5B**). The resulting MPs (W/O/W and S/O/W) exhibited similar EPO loading: 0.77 and 0.73 µg EPO per mg MP for W/O/W and S/O/W, respectively. Notably, W/O/W-fabricated MPs showed appreciable porosity and dimpling due to the formulation process, while S/O/W MPs had visually smoother surfaces with less porosity noted (**Figure 2A**). The formulations exhibited similar size, with a median diameter of ∼50 µm (**Supplementary Figure 5C-D**). We repeated longitudinal EDS to characterize whether protein loading changed MP oxidation dynamics in 25 mM H_2_O_2_, using bovine serum albumin (BSA) as a model for EPO due to its similar molecular weight and charge. These data suggest that S/O/W formulations oxidized faster than unloaded or W/O/W MPs, exhibiting complete dissolution after day 15 compared to day 30 (**Supplementary Figure 5E**). *In vitro* release of rhodamine-tagged BSA demonstrated sustained cargo release from both formulations over 110 days and that MP treatment with 10 mM H_2_O_2_ accelerated release (**Supplementary Figure 5F**). S/O/W MPs exhibited lower burst release than the W/O/W, perhaps related to the higher porosity of the W/O/W MPs.

**Figure 2:**
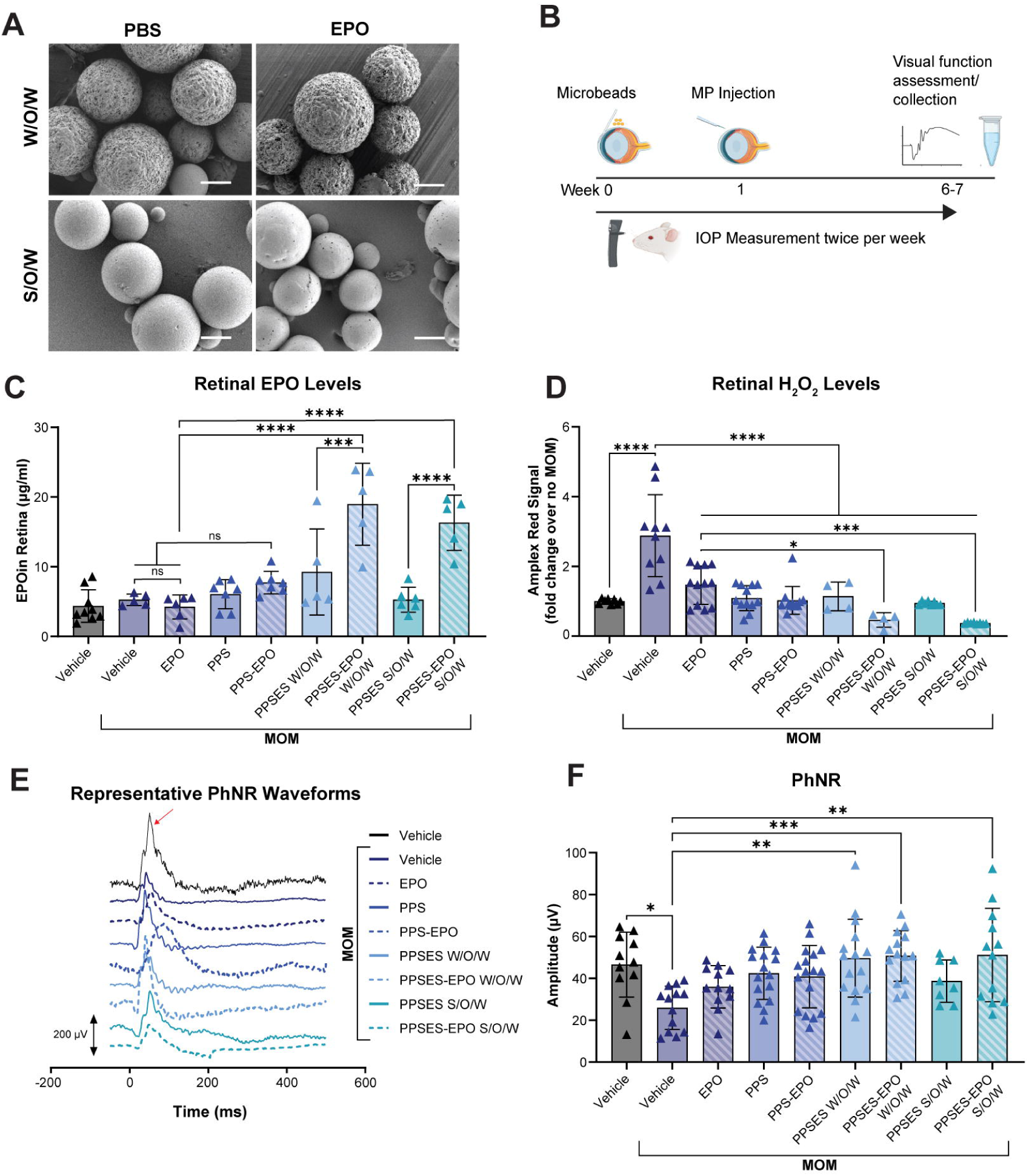
PPSES-EPO microparticles (MPs) protect the murine retina. **(A)** Representative scanning electron micrographs of PPSES MPs formulated by water-in-oil-in-water (W/O/W) and solid-in-oil-in-water (S/O/W) techniques. Scale bar = 60 pm. (B) Schematic of study design. **(C)** EPO in retina at endpoint (day 35-42 after MP injection) measured by ELISA. **(D)** *Ex vivo* retinal H,O_2_ levels measured by Amplex Red assay. (E) Representative photopic negative response (PhNR) waveforms with arrow indicating amplitude measurement. **(F)** Quantification of PhNR amplitude, ns = not significant, *p<0.05, **p<0.005, ***p<0.0005, ****p<0.0001. IOP = intraocular pressure; MOM = microbead occlusion model.

We tested our formulations in the well-characterized mouse microbead occlusion model (MOM) of glaucoma. In this model, IOP is elevated by injection of polystyrene microbeads into the anterior chamber of mouse eyes (*36*). Because PPS-EPO MPs previously exhibited promising results in our mouse model of blast-induced traumatic optic neuropathy (bITON) (*22*), another model of optic nerve degeneration, we treated with PPS-EPO MPs as a benchmark to compare with the effects of the new PPSES MP formulations. The MPs were injected (10 μg MP, ∼7.5 ng EPO) into the vitreous of the eye one week after induction of elevated IOP, and mice were then maintained for 6-7 weeks (**Figure 2B**). IOP levels were not affected by MP treatment (**Supplementary Figure 6A**). Both W/O/W and S/O/W PPSES-EPO MP formulations retained measurable amounts of EPO in the retina at study endpoint, which was not achieved with free EPO or PPS-EPO MPs (**Figure 2C**). PPSES-EPO MPs and PPSES-blank controls significantly reduced retinal H_2_O_2_ levels at a magnitude similar to or greater than PPS-based benchmark formulations (**Figure 2D**). Free EPO created an intermediate level of decrease in retinal H_2_O_2_ concentration and was significantly lower than vehicle-treated animals. Notably, both PPSES-EPO formulations significantly outperformed free EPO in retinal H_2_O_2_ scavenging.

Mice receiving vehicle (injection buffer) treatment with MOM had significantly decreased photopic negative response (PhNR) amplitudes at 6-7 weeks after IOP elevation, consistent with previous studies (*22*); however, treatment with PPSES-EPO MPs significantly improved the PhNR (**Figure 2E-F**). Assessment of bipolar cell function via the ERG b-wave demonstrated trending improvements with MP treatment, but did not reach statistical significance (**Supplementary Figure 6B-C**). To assess RGC axon transmission of visual information to the brain, we measured the visual evoked potential (VEP). Consistent with our previous studies (*7*, *32*), we detected a decrease in VEP N1 amplitude following MOM induction (**Figure 3A-B**). While all MP-EPO treated groups trended towards improvement, only S/O/W-formed PPSES MPs reached statistical significance over vehicle treatment. Notably, this formulation also significantly outperformed PPS, which we attribute to its higher stability (**Figure 3B**). To complement VEP results, optic nerve histology was assessed at the endpoint for all treatment groups (**Figure 3C-D**). Notably, free EPO did not maintain axon number, exhibiting total axon counts similar to treatment with vehicle (**Figure 3D**). However, all antioxidant MP formulations significantly preserved axons over free EPO treatment. Taken together, these findings demonstrate the capability for PPSES-based MP delivery of EPO to preserve neuronal functionality in the mouse MOM of glaucoma.

**Figure 3:**
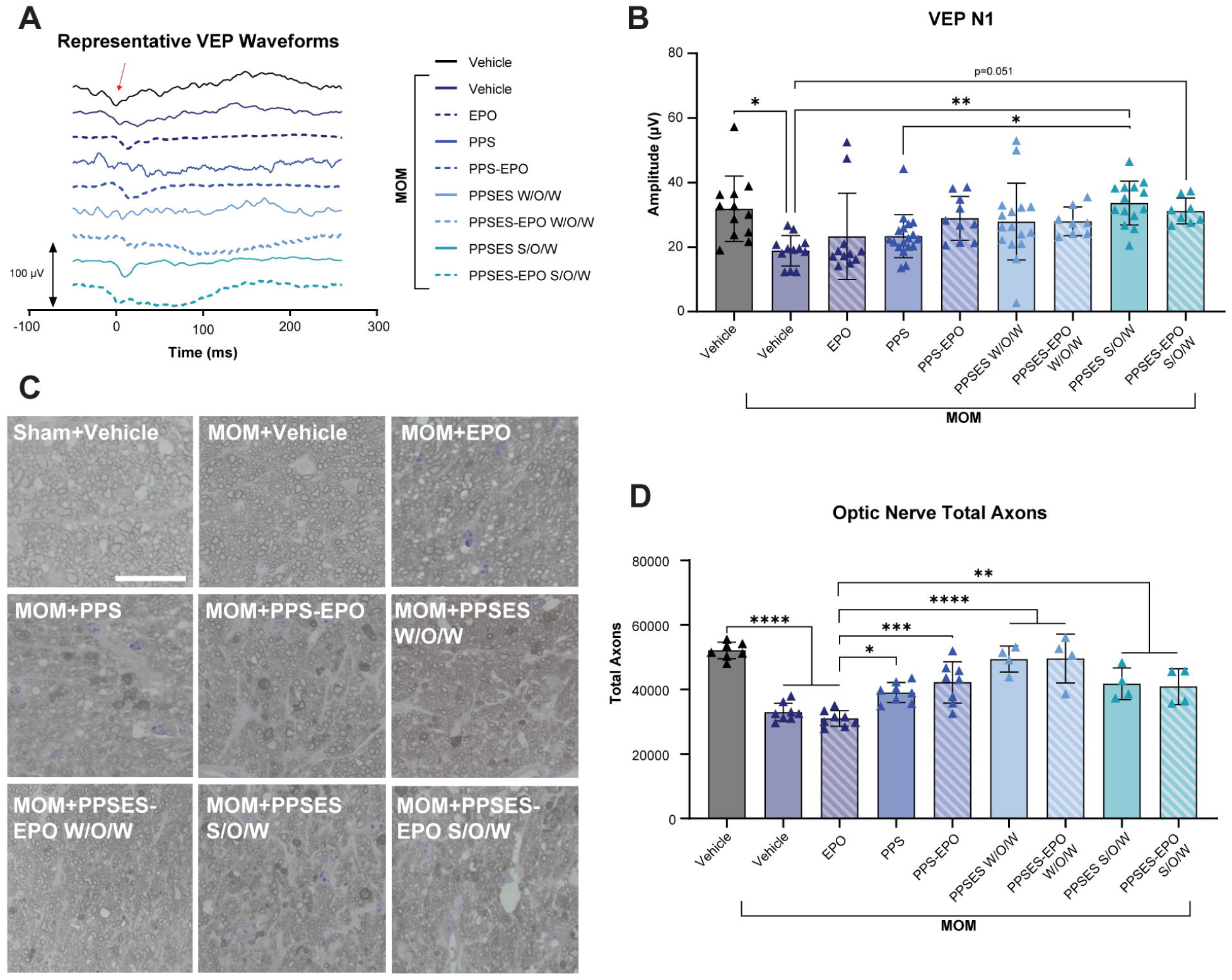
PPSES-EPO microparticles (MPs) protect the murine optic nerve. **(A)** Representative visual evoked potential (VEP) waveforms. Arrow indicates N1 wave. **(B)** Quantification ofVEPNl amplitude. (C) Representative micrographs of optic nerves. Scale bar = 20 pm. (D) Total axon counts of optic nerves. *p<0.05, **p<0.005, ***p<0.0005, ****p<0.0001. MOM = microbead occlusion model; S/O/W = solid-in-oil-in-water; W/O/W = water-in-oil-in-water.

### Optimization of PPSES-EPO MPs towards a NHP model of glaucoma

Our results in the mouse MOM supported pursuit of the S/O/W formulation for further testing in a human-relevant species. To increase the translatability of the system, we adjusted the formulation parameters to increase EPO loading (10.2 µg EPO/mg MP) and ensured that endotoxin levels were below the International Standards Organization (ISO) endotoxin safety limits for intraocular use (<0.5 EU/ml (*37*), **Supplementary Figure 7A**). The resulting PPSES-EPO MPs exhibited smooth morphology and the absence of porosity and dimpling exhibited in prior formulations (**Figure 4A**), while maintaining a similar median diameter of 56.0 µm (**Supplementary Figure 7B**). Using rhodamine-tagged EPO, it was observed that EPO was dispersed relatively homogeneously throughout the MP matrix (**Figure 4B**), and that protein release *in vitro* was accelerated in the presence of H_2_O_2_ (**Supplementary Figure 7C**). To further characterize the early transcriptional response to PPSES-EPO treatment, we performed RNA sequencing (**Supplementary Figure 8A-B**) 3 weeks after MOM induction, corresponding to 2 weeks after MP injection (**Figure 4C**). Differential expression analysis revealed that transcriptional profiles from the buffer-treated MOM animals and the PPSES-EPO-treated MOM retinas were inversely correlated, indicating that PPSES-EPO MP treatment broadly reversed ocular hypertension-induced gene expression changes (**Supplementary Figure 8C**). Consistent with this finding, gene set enrichment and pathway analyses demonstrated that biological processes associated with innate immune activation, inflammation, and oxidative stress were significantly induced in MOM retinas and partially normalized following PPSES-EPO MP treatment (**Figure 4D**). These effects were driven in part by altered expression of genes involved in glutathione metabolism (*Gsta3*, *Gpx3*), mitochondrial function and metabolic regulation (*Hmgcs2*, *Ucp2*), and innate immunity (*Lyz2*, *Ifit3*). Together, these findings demonstrate that sustained EPO delivery broadly counteracts transcriptional responses to ocular hypertension, including pathways associated with inflammation, oxidative stress, and metabolic dysfunction, providing a molecular basis for the functional and structural neuroprotection observed with PPSES-EPO MPs.

**Figure 4:**
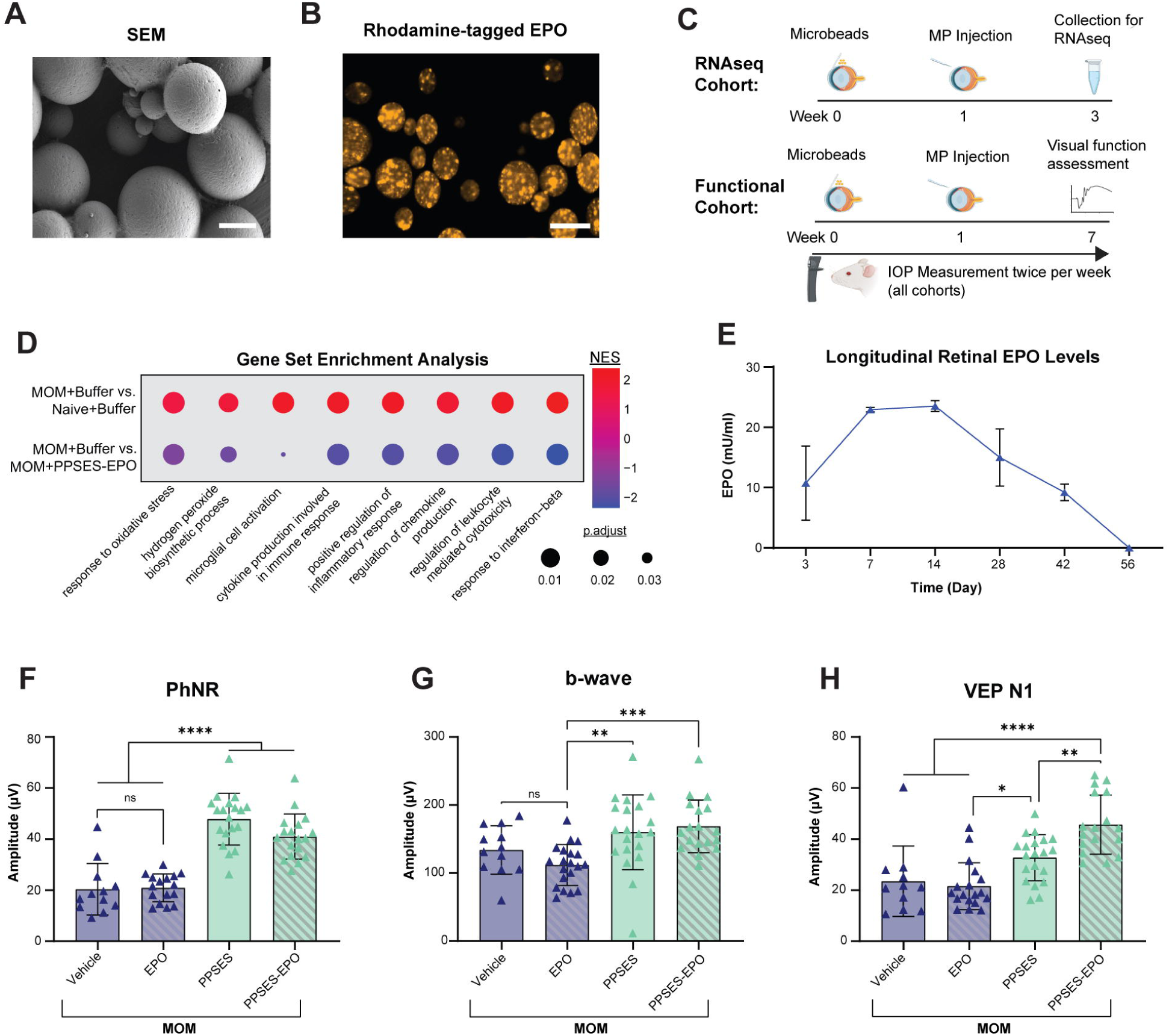
PPSES-EPO microparticles (MPs) optimized for translational application maintain efficacy in the mouse microbead occlusion model (MOM). (A) Scanning electron micrograph (SEM) of PPSES-EPO MPs. Scale bar = 50 pm. (B) Confocal microscopy demonstrating distribution of rhodamine-labeled EPO throughout the MPs. Scale bar =100 pm. **(C)** Schematic of study designs for RNA sequencing and functional assessment. (D) Gene set enrichment analysis of retinas with the indicated treatments 2 weeks after MP injection (3 weeks after initial MOM). **(E)** ELISA of retinas from PPSES-EPO MP-injected animals over time. (F-H) Electrophysiology 6 weeks after MP injection (7 weeks after initial MOM). (F) Quantification of photopic negative response (PhNR) amplitude. (G) Quantification of electroretinogram (ERG) b-wave amplitude. (H) Quantification of visual evoked potential (VEP)Nl-wave amplitude. *p<0.05, **p<0.005, ***p<0.0005, ****p<0.0001. IOP = intraocular pressure; NES = normalized enrichment score.

We next sought to determine the optimal dosing scheme for PPSES-EPO MP in NHPs based on longitudinal EPO release in the mouse MOM model. As before, MPs were injected 1 week after MOM induction (**Figure 4C**), and IOPs remained elevated throughout the study, with no difference in elevation among particle-injected groups (**Supplementary Figure 9A**). EPO ELISA on samples collected longitudinally demonstrated that EPO levels peaked at 7 and 14 days after injection and remained elevated out to 42 days post-injection, but were no longer measurable at day 56 (**Figure 4E**). Despite high local concentrations of EPO, no elevation in hematocrit was observed, indicating EPO remained in the eye (**Supplementary Figure 9B**). PhNR, ERG, and VEP tests at 7 weeks after initial MOM in PPSES-EPO MP-treated mice demonstrated excellent preservation of visual function, with significant improvement across measures for the PPSES-EPO MPs compared to vehicle and free EPO treatment (**Figure 4F-H, Supplementary Figure 9C-E**). Mice injected with optimized PPSES MPs, both with and without EPO loading, had significantly preserved PhNR (**Figure 4F**). PPSES-EPO also outperformed EPO and vehicle in preserving the ERG b-wave (**Figure 4G**). Finally, the VEP N1 amplitude was also significantly preserved with PPSES-EPO over all other treatment groups at week 7, with an intermediate effect of blank PPSES MPs (**Figure 4H**). Taken together, these results indicate strong neuroprotection by unloaded and EPO-loaded PPSES formulations.

These results supported testing of PPSES-EPO MPs in the NHP MOM (*38*). NHPs were injected intracamerally, bilaterally, with microbeads as needed until IOP increased (**Supplementary Figure 10**). After 6 weeks of elevated IOP, NHPs were intravitreally injected with buffer, PPSES MPs, or PPSES-EPO MPs every 5 weeks until collection at 45 weeks (**Figure 5A**). One NHP received injections of MPs loaded with rhodamine-tagged EPO for detection of MP biodistribution. We detected EPO-loaded MPs on the surface of the retina 5 weeks after the last injection, with additional EPO signal visible throughout the retina, indicating diffusion from the MPs (**Figure 5B**). Similar to the murine model, NHPs exhibited normal hematocrit (42±2%), and we did not detect an increase in serum EPO levels. Altogether, these data indicate excellent retention of EPO within the eye and lack of systemic effects using this treatment approach.

**Figure 5:**
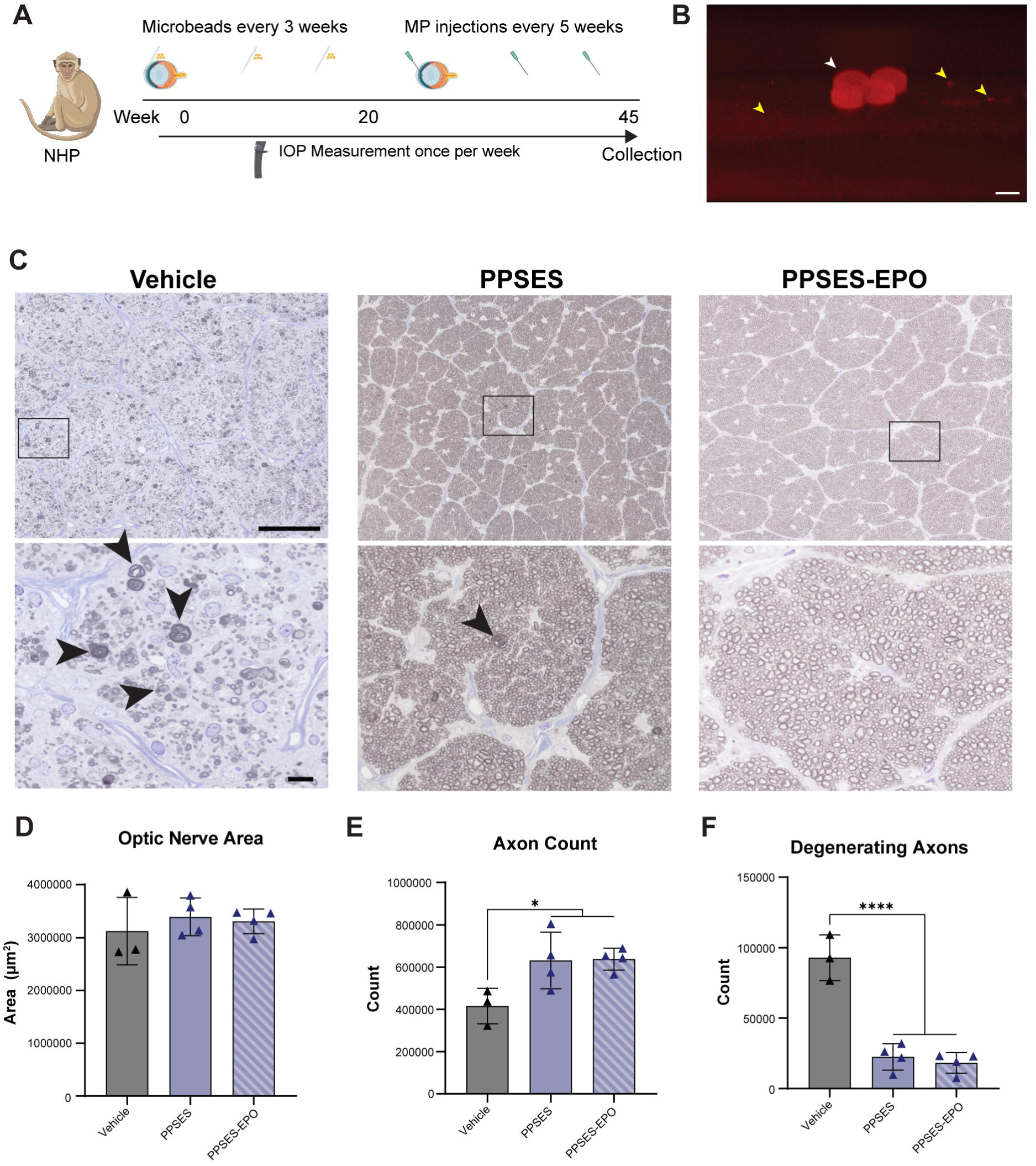
Efficacy of PPSES-EPO microparticles (MPs) in the non-human primate (NHP) microbead occlusion model (MOM). **(A)** Study schematic. **(B)** Confocal microscopy demonstrating PPSES-EPO MPs loaded with rhodamine-labeled EPO deposited on the NHP retina. White arrow indicates intact MPs, and yellow arrows indicate diffused EPO. Scale bar = 50 pm. (C) Representative micrographs of NHP optic nerve cross sections. Arrows in zoom boxes indicate degenerative axons. Top scale bar = 25 pm; zoom scale bar = 10 pm. **(D)** Quantification of optic nerve area. **(E)** Total axon counts and **(F)** degenerative axon profiles from optic nerve cross sections. *p<0.05; ****p<0.0001. IOP = intraocular pressure.

Our previous work showed an increase in multiple pro-inflammatory cytokines in the glaucomatous NHP aqueous humor (*38*); therefore, we measured IFN-γ levels in a subset of animals. In PPSES and PPSES-EPO treated animals, we detected no increase in IFN-γ (**Supplementary Table 1**). Neuroprotection was assessed through histological evaluation of the optic nerve (**Figure 5C**). We detected consistent optic nerve cross area in all treatment groups (**Figure 5D**). Both PPSES MP formulations (unloaded and EPO-loaded) significantly preserved total axon counts (**Figure 5E**) by robustly preventing axon degeneration (**Figure 5F**). Therefore, by these metrics, MPs alone were as effective as EPO-loaded MPs in protecting the optic nerve axons in a NHP model of glaucoma. The efficacy of the empty,H2O2-scavenging PPSES MPs emphasizes the important role of elevated ROS in the pathogenesis of glaucoma.

## DISCUSSION

Over 95 million people worldwide are affected by glaucoma (*39*), and although current treatments can effectively lower IOP in some patients, many patients progress to blindness despite IOP management, highlighting the need for new neuroprotective strategies. Here, we optimized an injectable, sustained-release MP system with comprehensive antioxidant activity via [1] direct ROS scavenging by PPSES and [2] NRF2 activation by EPO. Oxidative stress plays a key role in glaucoma pathogenesis (*2–5*, *40*, *41*); therefore, we posited that the polysulfide chemistry of PPSES would contribute directly to therapeutic benefit through ROS scavenging (particularly H_2_O_2_, hypochlorite, and peroxynitrite) (*21*). Additionally, there is a large body of preclinical evidence on the neuroprotective effect of EPO in the optic nerve, retina, and brain (*25–28*, *42*, *43*) due to its activation of antioxidant enzyme production (*32*). There are also ongoing clinical trials of EPO as a therapy for spinal shock (NCT00561067), critical trauma (NCT04588311), and Parkinson’s disease (NCT01010802). As glaucoma is a progressive, lifelong disease, it necessitates sustained therapeutic activity to minimize the number of intravitreal injections and systemic side effects while maximizing efficacy. While sustained release of EPO has already been accomplished by our group and others using PLGA (*29*, *30*, *32*, *44*) and PPS (*22*), PPSES MPs preserve the ROS scavenging of PPS while providing increased size and stability seen in gold-standard PLGA systems. Together, our findings establish PPSES MPs as a sustained antioxidant delivery platform with robust neuroprotective efficacy across murine and primate models of glaucoma.

We successfully optimized PPSES monomer composition (50:50 PS:ES mol %) for sufficient crystallinity to yield stable drug-releasing MPs at 10-100 µm (**Figure 1 and Supplementary Figure 1 and 2**). The broad crystallization and melting DSC peaks (**Figure 1A** and **Supplementary Figure 2C**) were likely caused by the nonstochastic integration of ES monomer into the polymer chains due to the difference in reactivity of the two monomers;(*45*, *46*) indeed, complex lamellae were confirmed using SSA (**Supplementary Figure 2D**). As expected, the end temperature of the melt (T_m,end_) was higher at 50 mol% ES than 40 mol% ES (82.3°C vs. 55.1°C, respectively), which suggests this polymer is more crystalline and exhibits greater physical integrity at physiological temperature (37°C). Taken together, our data demonstrate that detectable crystallinity is required to generate MPs in the 10-100 µm diameter range, and to our knowledge, this is the first application of PPSES copolymers for large MP fabrication.

Loading proteins into organic matrices is challenging due to poor encapsulation and structural degradation. W/O/W formulations are commonly used due to their simplicity but suffer from low cargo loading and burst release profiles (*47*), prompting us to pursue a S/O/W technique to address these issues (*48*, *49*). We first compared W/O/W and S/O/W PPSES-EPO MPs, which exhibited similar size and EPO loading (**Figure 2A**). The sustained release of S/O/W PPSES-EPO MPs in PBS was comparable to PLGA-EPO MPs (*31*) and decreased the burst in the early time points compared to W/O/W (**Supplementary Figure 5F**). The slower burst release could be due to protein interactions with the DEAE-dextran, since electrostatic interactions can slow release of proteins (*50–52*). H_2_O_2_ accelerated protein release (**Supplementary Figure 5F and Supplementary Figure 7C**), corroborating the ROS-responsiveness of the polymer via thioether oxidation, which was also demonstrated by EDS (**Figure 1D-E** and **Supplementary Figure 3 and 4**). The rate of MP breakdown was dependent on the dose of H_2_O_2_ and correlated with increased oxygen percentage, suggesting that MP erosion is dictated by level of oxidation.

We proceeded to confirm sustained EPO release and antioxidative effect *in vivo*. Because intravitreal injections are commonplace at intervals of 4-8 weeks for chronic vascular diseases (*53*), we targeted sustained *in vivo* release of EPO for at least 4 weeks. Indeed, PPSES delivery of EPO maintained elevated EPO levels in the retina for 6 weeks (**Figure 2C and 4E**), while eyes treated with free EPO or PPS-EPO MPs did not have measurable EPO levels at this time point (**Figure 2C**). The improved EPO delivery for PPSES compared to PPS is likely due to the increased size, retention, and stability of PPSES. Mice (**Supplementary Figure 9B**) and NHPs locally injected with PPSES-EPO MPs did not exhibit elevated hematocrit, a notable side effect of systemic EPO delivery, indicating safety of the system. ROS scavenging by PPSES and PPSES-EPO MPs were also confirmed in the murine retina, where all PPSES MP treatments robustly reduced H_2_O_2_ signal 6 weeks after MP injection (7 weeks after initial MOM, **Figure 2D**). Notably, the addition of EPO to the MPs resulted in a trend toward further reduction in H_2_O_2_ signal compared to unloaded PPSES MPs, although this difference was not statistically significant. These findings are consistent with our previous study demonstrating that retinal oxidative stress in the mouse MOM peaks at early time points (1–2 weeks post-IOP elevation), coinciding with activation of the NRF2/ARE pathway (*7*, *32*, *54*). These results informed the treatment schedule in the NHPs, in which optimized PPSES-EPO MPs were injected every 5 weeks. MPs were detected on the retina 5 weeks after the last injection, along with diffused EPO throughout the tissue (**Figure 5B**), indicating that this retreatment paradigm would likely be effective in patients. Future studies are needed to determine if the therapeutic effect would be sustained with longer injection intervals. Together, these findings demonstrate that PPSES MPs provide sustained local EPO delivery and antioxidant activity while limiting systemic EPO exposure, supporting their potential for long-term treatment of glaucoma.

Potent ROS scavenging and prolonged EPO release resulted in significant neuroprotection and functional preservation. In the murine MOM, we detected significant improvements in bipolar cell, RGC, and optic nerve function at 6 weeks after MP injection (7 weeks after initial MOM) by electrophysiology (**Figure 2E-F**, **3A-B**, **4F-H**; **Supplementary Figure 9C-E)**. Murine optic nerve axons were also significantly preserved at this time point (**Figure 3C-D**). Notably, retinal EPO levels declined over time and were no longer measurable at 8 weeks after MP injection (**Figure 4E), highlighting** the importance of establishing an appropriate retreatment interval to maintain sustained therapeutic activity. Consistent with these neuroprotective effects, early transcriptomic analysis demonstrated that PPSES-EPO treatment attenuated ocular hypertension-associated transcriptional signatures related to oxidative stress, mitochondrial dysfunction, and inflammatory signaling (**Figure 4D**). Together with our findings at later time points (**Figure 2C-F**, **3A-D**, **4F-H**), these results suggest that PPSES-EPO modulates molecular pathways associated with retinal injury early after treatment, potentially contributing to the sustained functional protection observed.

With repeated dosing in the NHP, the optic nerve axons were significantly preserved (**Figure 5C-F**). Interestingly, the results of this study show that blank PPSES MPs were statistically similar to PPSES-EPO MPs across many of the neuroprotective metrics tested. This is possibly due to saturation in these metrics, and future studies measuring active axon transport or visual function in the NHP may identify differences not detected with the assays included in this study. The robust neuroprotection of PPSES suggests that H_2_O_2_, hypochlorite, and/or peroxynitrite are key ROS in glaucoma pathogenesis. It also suggests that decreasing these ROS is sufficient to suppress inflammation, and that glaucomatous axon degeneration is at least partially dependent on ROS levels. Notably, clinical evidence for dietary antioxidants to mitigate glaucoma pathogenesis is mixed, with many studies failing to demonstrate significant therapeutic benefit (*55*). This suggests that local ROS scavenging could represent a superior strategy. Since oxidative stress is a common pathway in many neurodegenerations, our data indicate that PPSES represents a promising new biomaterial for antioxidant activity and sustained cargo delivery in glaucoma, and could be extended to other neurodegenerative disorders.

## CONCLUSIONS

Overall, PPSES-EPO MPs are a promising candidate for intravitreal treatment of glaucoma. Careful tuning of polymer characteristics was essential to achieve sustained effects in small and large animal models. This system exhibits potent ROS scavenging and neuroprotection and has been adapted to a translationally-relevant NHP model. Rescue of retina and optic nerve is extremely challenging clinically, necessitating innovative therapies like PPSES-EPO MPs.

## MATERIALS AND METHODS

### Materials

Unless otherwise stated, all materials were purchased from Sigma Aldrich or ThermoFisher and used as is. All assay kits were performed according to manufacturer’s instructions unless otherwise stated.

### Microparticle synthesis and characterization

#### Polymer synthesis

Propylene sulfide (PS) was distilled to remove inhibitor and stored at −20C until use. 2,2’-(ethylenedioxy)diethanethiol was used as initiator. Initiator (0.17 mmol), 1,8-diazabicyclo(5.4.0)undec-7-ene (DBU, 0.67 mmol), and tributylphosphine (TBP, 1.7 mmol) were mixed in 10 mL degassed tetrahydrofuran (THF) at room temperature. PS and ethylene sulfide (ES) were combined in a single vial at molar ratios varying from 100:0 to 50:50 PS:ES, with a total monomer amount of 36.9 mmol. The monomer mixture was added to the reaction in 8 equal portions, with 15 minutes between additions, followed by an additional 45 minutes of reaction time. 1-iodopropane (0.50 mmol) was added to end cap the polymer and reacted for 1-3 hours. THF was removed by rotary evaporation, crude product was redissolved in dichloromethane DCM and extracted 3x against brine. DCM was removed by rotary evaporation, and the concentrated polymer was precipitated 3x in MeOH. The purified polymer was redissolved in ∼40 ml DCM, and insoluble fragments were removed via filtration through coarse filter paper. Excess DCM was removed by rotary evaporation and dried in a vacuum oven. Samples were analyzed by NMR (Bruker) at ∼1% (w/v) in deuterated chloroform (CDCl_3_). Conversion of the PS and ES monomers was confirmed by integrating the ^1^H NMR (400 MHz),δ= 2.9, 2, CH_2_; 2.8, 4, CH_2_; 2.6, 1, CH; 1.4, 3, CH_3_; 3.6, CH_2_, 3.7, CH_2_. The relative molecular weight of the series was determined via GPC as described below.

#### GPC

The molecular weights and polydispersity indices were measured using gel permeation chromatography (GPC). Analysis was performed using a EcoSEC Elite GPC System with LenS MALS and refractive index (RI) detectors (Tosoh Biosciences, Japan) comprising of a TSKgel Alpha-M 13 μm column. The mobile phase consisted of DMF with 0.01 M LiBr used at a flow rate of 0.6 mL/min and a temperature of 50 °C. Polymers were dissolved at 2 mg/ml and filtered with a 0.22 um PTFE filter before analysis. Polymer refractive index increment (dn/dc) values were calculated using a digital refractometer (Anton Paar) using polymers dissolved in the same mobile phase at concentrations of 0.5, 1, 2.5, 5, 7.5 and 10 mg/mL. GPC calibration was performed using polymethylmethacrylate (PMMA) standards of known molecular weight and dn/dc.

#### DSC

Approximately 10 mg polymer was weighed into an aluminum Tzero pan, crimped, and loaded into the calorimeter (TA Instruments). Air was used as a reference sample. The sample was stabilized at 140°C for 3 minutes to erase thermal history. Next, the sample was cooled at 20°C/min to −80°C to promote crystallization and glass transition. After 3 minutes to stabilize, the sample was heated to 140°C at 20°C/min to capture melting events.

#### SSA

PS_50_-co-ES_50_ was prepared and loaded into the calorimeter as above. The sample was stabilized at 150°C for 6 minutes to erase thermal history. For all subsequent temperature changes, the rate was held constant at 5°C/min. The sample was equilibrated at −50°C to promote crystallization and glass transition. To achieve thermal fractionation, a temperature cycle was established between −50°C to 70°C, and the maximum temperature was decreased by 5°C each cycle (i.e. 65, 60, 55°C, etc.) to −45°C. Finally, the polymer was heated to 150°C to capture the melting of each established lamellae.

#### XRD

Polymer samples were deposited onto a glass sample holder and placed in the vacuum chamber of the XRD instrument scanner (Rigaku). Reflections were captured at 2θ= 10-30 at 9°/min with a step size of 0.1°. Data was smoothed and normalized to peak intensity for sample comparison. The X_c_% was determined by measuring the AUC at 2θ= 24° and comparing to the total AUC.

#### MP fabrication

Polymers were dissolved in organic phase comprising of DCM or 50:50 DCM:CHCl_3_ at 50 mg/ml w/v. Oil phase was added to water phase comprising 10-15 mg/ml PVA (w/v), with or without 50 mg/ml NaCl (w/v). The O/W emulsion was prepared either via homogenization or stirring at various speeds to control MP size. Polymer/oil droplets were hardened to MPs either via rotary evaporation (10-60 min) or gentle stirring in a hardening solution composed of 1mg/ml PVA (w/v) with or without 50 mg/ml NaCl (/wv) and 0.5% Tween-20 (v/v) (10-60 min). MPs were centrifuged at 300xg for 3 minutes and rinsed with DI water 1-2x. Samples were resuspended in water or a lyoprotectant solution of 6.67 mM sucrose and 0.5% Tween-20, snap frozen in liquid nitrogen, and lyophilized.

#### SEM

Samples were suspended in DI water and deposited onto steel stubs coated with copper tape (Electron Microscopy Sciences). Dextran NPs were suspended in DCM instead of water to maintain morphology. After air drying, the sample was sputter coated with gold. MPs were imaged on a Nikon SEM with Gemini column.

#### Oxidation-dependent degradation of MPs

MPs were resuspended at 10 mg/ml in 0.25% Pluronic F-127 + 0.25% carboxymethylcellulose. H_2_O_2_ was added for final concentrations of 0, 10, 25, 50, and 100 mM H_2_O_2_ and 2.5 mg/ml MP. Samples were agitated at 37°C for the duration of the experiment. At pre-determined time points, 60 μl samples were collected, resuspended in water, and deposited onto SEM stubs for morphological and elemental analysis. Stock tubes were centrifuged at 300xg, and supernatant was replaced with fresh H_2_O_2_ at each collection time point. 0.5% Tween-20 was incorporated into all solutions to prevent MP adherence to tube and pipette tip walls. Morphology was visualized via SEM as described above.

#### EDS

At each time point, samples were analyzed for elemental composition via EDS on the Nikon SEM. Applied voltage and current were 10 kV and 1 nA, respectively, and collection time was 20 seconds. Weight percentage of the three major elements, C, O, and S, were calculated from the resulting spectra. MPs were categorized as“small” (<50μm) and“large” (>50μm), and five spectra per size per sample were collected.

#### MP cytotoxicity

ARPE-19 cells were seeded at 10,000 cells/well into a 96-well plate in serum-free DMEM/F12. Cells were allowed to adhere for 24hr. Once cells adhered, media was replaced with 100 μl media containing MP suspensions at 0.1-2 mg/ml. After 24 hours of treatment with MPs, cell viability was measured using a Cell TiterGlo kit (Promega) according to the manufacturer’s protocol.

### Fabrication of drug-loaded MPs

#### W/O/W emulsion for EPO MPs

PPSES-EPO W/O/W MPs were formulated with the following conditions: 50 μl EPO-R76E (450 μg/ml) was added to 1 ml PPSES in DCM (11.25 mg/ml). The W/O emulsion was homogenized at 20,000 rpm for 30 seconds, then added dropwise to 6 ml 1% PVA +10% NaCl. The W/O/W emulsion was stirred at 800 rpm for 60 seconds and hardened in 200 ml 0.1% PVA+ 10% NaCl for 4 hours on ice. The MPs were collected and rinsed 1x with water. The hardened MPs were resuspended in 1 ml 6.67 mM sucrose + 0.5% Tween-20, frozen in liquid nitrogen, and lyophilized. Control MPs were formulated with 50 μl PBS as the internal W phase. EPO encapsulation was determined by degrading the MPs in 1000 mM H_2_O_2_ for 48 hours and measuring EPO concentration by ELISA.

#### S/O/W emulsion for EPO MPs

PPSES-EPO S/O/W MPs: 100 μg EPO (Prospec) was reconstituted in ∼200μl DI water, transferred to a PCR tube, and lyophilized. The protein was reconstituted in 10 μl diethylaminoethyl (DEAE)-dextran (70,000 Da, 6 mg/ml) and incubated for 5 minutes on ice. 10 μl dextran (40,000 Da, 34 mg/ml) was added, followed by 20 μl PEG (8000 Da, 160 mg/ml). The suspension was vortexed at max speed for 45 seconds and snap-frozen in liquid nitrogen for lyophilization. The lyophilized NPs were washed 3x with DCM to remove PEG, suspended in 200 μl PPSES (50 mg/ml in DCM), and added dropwise to 6 ml 1% PVA. The sample was stirred for 60 seconds at 1000 rpm, rinsed 1x with DI water, and resuspended in 0.5% Tween-20 + 6.67 mM sucrose. MPs were flash-frozen in liquid nitrogen and lyophilized. Control MPs were formulated with blank DEAE-dextran NPs. EPO loading was determined by dissolving MPs in DCM, collecting dextran MPs by centrifugation, and rinsing twice more with DCM before lyophilization. Dextran/EPO pellet was dissolved in DI water, and EPO concentration was determined by ELISA (Boster Bio).

Formulation adjustments for NHP PPSES-EPO S/O/W MPs: For preparation of “low-endotoxin” MPs, the following sterilization protocol was followed: all aqueous solutions were prepared in sterile DEPC-treated water and sterile filtered prior to use. All glassware and stir bars were soaked in freshly-prepared 0.2 M NaOH overnight, rinsed twice with sterile saline, rinsed twice with sterile DEPC-treated water, and allowed to dry in a laminar flow hood. The samples were exclusively handled in laminar flow hoods except during the use and evaporation of volatile solvents DCM and CHCl_3_. Commercial EPO (Antibodies Online) was reconstituted at 100 µg/ml in 1X PBS and concentrated via centrifugation at 4°C through a 10kDa spin filter to ∼20 mg/ml. EPO was aliquoted, and final concentration was determined by ELISA. 18 μl DEAE-dextran (4.69 mg/ml) was added to 2 μl EPO (562 μg/μl) and incubated for 30 minutes on ice. 10 μl dextran (36.59 mg/ml) was added, followed by 15 μl PEG8000 (240mg/ml). The solution was vortexed at max speed for 60 seconds, incubated on ice for 10 minutes, and frozen in liquid nitrogen for lyophilization. The lyophilized NPs were washed 3x with DCM to remove PEG, suspended in 200 μl PPSES (50 mg/ml, CHCl_3_/DCM), and added dropwise to 1ml 1% PVA + 5% NaCl. MP droplets were hardened for 15 minutes in 200 ml 0.1% PVA + 5%NaCl + 0.5% Tween-20. Hardened MPs were collected and rinsed twice with DI water, resuspended in DI water, snap frozen, and lyophilized. Control MPs were formulated with blank DEAE-dextran NPs. EPO encapsulation was determined by measuring EPO concentration in each wash step by ELISA (Boster Bio). Control samples were prepared by suspending dextran-EPO MPs in a DCM/CHCl_3_ oil phase, emulsified, and processed as MPs above. Encapsulation efficiency in EPO MPs was determined by comparing EPO concentration (ELISA) in the supernatants of control and MP samples.

#### Endotoxin measurement

MPs were resuspended in injection buffer, and 50 μl were plated in a 96-well plate. Endotoxin was measured using the Pierce Chromogenic Endotoxin Quant Kit (based on limulus amebocyte lysate (LAL) according to manufacturer’s directions with the following adjustment: after stop solution was added and absorbance was measured, a second measurement was taken on reaction supernatants. This helped to correct for absorbance interference from sedimented MPs in the wells.

### Murine MOM and treatments

#### Mice

C57Bl/6 J mice (Jackson Laboratories) were group-housed, maintained on a 12-h light-dark cycle, and provided food and water *ad libitum*. An equal distribution of 2-3 month old male and female mice were used. All animal studies were performed according to an institutionally approved protocol within AALAC accredited laboratories.

#### Murine MOM

IOP was bilaterally elevated using the well-characterized MOM. 15-µm diameter FluoSpheres polystyrene microbeads (Spherotech) were injected into the anterior chamber of anesthetized mice as previously described (*36*, *56*). Briefly, 1.5 mm outer diameter/1.12 mm inner diameter filamented capillary tubes (World Precision Instruments) were pulled using a P-97 horizontal puller (Sutter Instrument Company), and the resulting needles were broken using forceps to an inner diameter of ∼100 µm. Microbeads were loaded and injected using a microinjection pump (World Precision Instruments). Mice were anesthetized with isoflurane and dilated using topical 1% tropicamide ophthalmic solution (Patterson Veterinary), and 2 µl (∼2,000 microbeads) were injected. The needle was maintained in the injection site for 20 seconds before retraction to reduce microbead efflux. Mice were given topical 0.3% tobramycin ophthalmic solution (Patterson Veterinary) following injection.

#### Murine IOP measurements

IOP was measured immediately prior to microbead injection and biweekly thereafter using the Icare TonoLab rebound tonometer (Colonial Medical Supply) as previously described (*7*, *32*, *36*, *42*, *54*). Mice were anesthetized using isoflurane, and 10 measurements were acquired from each eye within 2 minutes of induction of anesthesia.

#### Murine MP treatment

For all murine studies, intravitreal injection of treatments occurred 1 week after the first microbead administration. All injections were 1 µl, and for MP groups, all animals received 10 µg MP. Microbeads were re-administered every 3-4 weeks to maintain IOP elevation. **Supplementary Tables 2-4** summarize the sample sizes, treatment groups, and number of animals for the three murine studies presented here. For hematocrit measurement, blood was collected from the tail vein, and hematocrit was measured by capillary centrifugation.

### Murine in vivo electrophysiology

ERG and VEP measurements were performed as previously described (*7*, *57–60*). Mice were dark-adapted overnight before being anesthetized for electrode placement at the cornea, snout, and back of the head (visual cortex). For VEPs, mice were exposed to 50 flashes of 1 Hz of 0.05 cd.s/m^2^ white light. For ERGs, mice were exposed to flashes of 1 Hz of 1 cd.s/m^2^ white light. For the photopic negative ERGs (PhNR), mice were exposed to 20 flashes of white light on a green background. Testing was performed in order of dimmest to brightest light exposure such that animals received PhNR at the end of the testing session.

#### Amplex Red assay

Freshly harvested retinas were placed on ice until Amplex Red assay according to the manufacturer’s protocol with the following adaptations: Assay reagent was prepared by combining 11.82 ml Kreb’s buffer (pH 7.4), 120 μl HRP, and 60 μl Amplex Red dye. Retinas were suspended in 500 μl reagent and homogenized by hand. Samples were incubated at 37°C, centrifuged at max speed for 5 minutes, and the supernatant was plated in duplicate. Absorbance was measured at 570 nm.

#### Retinal EPO ELISA

For ELISA, retinas were flash frozen until analysis. Samples were sonicated, and supernatant was diluted in ELISA buffer (R&D Systems) for analysis according to manufacturer’s directions.

#### Optic nerve histology

Optic nerves were post-fixed in glutaraldehyde followed by Resin 812 embedding and Araldite 502 (Electron Microscopy Sciences) according to previously published protocols (*42*, *57*, *61*). 1 µm sections were stained with 1% paraphenylenediamine and 1% toluidine blue and were imaged on a Nikon Eclipse Ni-E microscope using 100x oil immersion objective (Nikon Instruments). The optic nerves were montaged into a 5 x 5 image using the Nikon Elements software to scan a large image. Healthy and degenerative axons were counted manually in ImageJ (Counting Array and Better Cell Counter plugins). A grid was used to avoid bias, by always counting in the same squares, using a cross configuration. 20% of the optic nerve cross-sectional area was counted and the total was multiplied by five to estimate total and degenerating axons within the nerve.

#### Bulk RNA sequencing

Retinas were processed under RNase-free conditions and homogenized in lysis buffer by sonication prior to RNA purification. Total RNA was isolated using the RNeasy Plus Mini Kit (Qiagen) according to the manufacturer’s instructions, including genomic DNA removal using gDNA eliminator columns. RNA quality control, library preparation, and sequencing were performed by the Vanderbilt Technologies for Advanced Genomics (VANTAGE) core. Libraries were generated using a total RNA rRNA-depletion workflow and sequenced on an Illumina NovaSeq X Plus platform to generate paired-end 150-bp reads with a target depth of ∼50 million reads per sample. Demultiplexed FASTQ files were processed using a standard RNA-seq pipeline including read quality assessment, adapter trimming as needed, alignment to the mouse reference genome, and generation of gene-level count matrices.

Raw read counts per gene were extracted using HTSeq-count (v0.11.2). Differential gene expression analysis was performed using the DESeq2 package (v1.38.3), with normalization by the median-of-ratios method. Genes with fewer than 15 counts across all samples were excluded from analysis. Multiple testing correction was performed using the Benjamini–Hochberg method to control the false discovery rate (FDR), with adjusted P < 0.05 considered statistically significant. Significantly perturbed pathways between groups were identified using clusterProfiler (*62*).

### NHP MOM

#### Primates

Adult male and female Bolivian squirrel monkeys (*Saimiri boliviensis*, MD Anderson), aged 2 to 4 years, were used in this study. All animals were free of any history of diabetes, head injury, or ocular trauma. They were socially housed, provided with environmental enrichment, and maintained on a 12-hour light/dark cycle. Animals had *ad libitum* access to water and were fed a balanced diet consisting of standard primate chow, biscuits, fresh vegetables, fruits, and other nutritional supplements. All animal studies were performed according to an institutionally approved protocol within AALAC accredited laboratories.

#### NHP MOM

Animals were sedated with midazolam (Avet Pharmaceuticals) and subsequently placed under general anesthesia with isoflurane. Under sterile conditions, bilateral intracameral injections (40 µl/eye) of 45 µm fluorescent carboxyl magnetic particles suspended in 2% hydroxypropyl methylcellulose (HPMC) in PBS were administered. Injections were repeated every 3 weeks until sustained elevation of IOP was achieved. Standard post-procedural monitoring was conducted, and NHPs were returned to social housing once fully ambulatory.

#### Awake IOP measurement

IOP was measured weekly in awake animals using a custom-designed restraint tube and positive-reinforcement training protocol. Briefly, NHPs were gradually conditioned to tolerate hand-catching, placement into the restraint apparatus, and tonometry using reward-based behavioral training. Habituation sessions were conducted Monday through Friday and included desensitization to the handling procedures, the restraint tube, and the use and sounds of the rebound tonometer (TonoPen XL; Medtronic Solan). Once acclimated, IOP measurements were obtained following topical instillation of 0.5% proparacaine hydrochloride ophthalmic drops (Alcon Laboratories, Inc.). Fifteen readings per eye were recorded per session and averaged for analysis.

#### Intravitreal injections

Intravitreal injections were performed under sedation with midazolam, followed by maintenance of general anesthesia with isoflurane. Using sterile technique and a 29-gauge needle, 20 µl of the assigned treatment. All injections were performed bilaterally and repeated every 5 weeks following the initial administration. **Supplementary Table 5** summarizes the treatments and sample sizes for NHP treatments. Animals were monitored post-procedure until fully ambulatory, and all treatments were well tolerated without injection-related complications.

#### Aqueous humor and blood sampling

During the same anesthetic event as the magnetic particle injections, aqueous humor and blood samples were collected for analysis. Aqueous humor (10-20 µl per eye) was aspirated from the anterior chamber using a 30-gauge × 5/16-inch insulin syringe under sterile conditions. Hematocrit samples were obtained via venipuncture of the saphenous vein using a 25-gauge needle. All procedures were performed under general anesthesia, and animals were monitored continuously throughout the sampling process.

#### Cytokine and EPO measurement

EPO levels were quantified from serum samples by ELISA, and 7 additional growth factors and cytokines were measured from aqueous humor samples using the Human Cytokine/Chemokine Magnetic Bead Panel A (Millipore). Both assays were performed according to the manufacturer’s instructions.

#### Optic nerve histology

Following enucleation, optic nerve segments approximately 1 mm in length were carefully dissected immediately posterior to the globe. Tissues were post-fixed in 2% glutaraldehyde for 2 hours to preserve ultrastructural integrity. After fixation, nerves were embedded in Epon resin and sectioned at 1-2 µm thickness for morphological evaluation using standard protocols. Sections were stained with 1% paraphenylenediamine and 1% toluidine blue. High-resolution images of optic nerve cross-sections were acquired using a Nikon Eclipse Ni-E microscope equipped with a DS-Ri2 color digital camera. Images were obtained with a 40× oil-immersion apochromatic objective under brightfield or DIC optics. Sequential fields were captured and automatically stitched into montages using NIS-Elements software to encompass the entire nerve profile at high spatial resolution. Pixel calibration was set to 0.18 µm/pixel for 40× images, based on manufacturer specifications for the DS-Ri2 camera and 1.0× adapter.

#### Axon quantification

Optic nerve cross-sections were analyzed in Fiji/ImageJ (NIH) using a custom plugin-ready macro that automated 8-bit conversion, rolling-ball background subtraction (25 px), global thresholding, binary conversion, and optional watershed separation to ensure consistent segmentation across all images. For intact axon counts, Otsu thresholding was applied with size and circularity filters of 0.2-20 µm² and 0.30-1.00, respectively, which reliably isolated individual axonal profiles while excluding debris and merged clusters. For degenerating profiles, Moments thresholding was applied with size and circularity filters of 0.25-10 µm² and 0.00-0.75, optimized to detect dark, vacuolated, or irregular axons. Degenerating profiles were morphologically defined by the presence of axoplasmic swelling or vacuolization, myelin sheath fragmentation or collapse, condensed axoplasmic material, and loss of circular contour; myelin ovoids and axonal fragments contiguous with disrupted fibers were also included. Morphometric features (area, circularity, aspect ratio, roundness, and solidity) were extracted for each axon.

### Statistics

All data is represented as mean +/- standard deviation unless otherwise stated. Data was analyzed by one-way ANOVA with Tukey’s post-hoc analysis unless otherwise stated. EDS oxidation data was evaluated by two-way ANOVA with Šídák’s multiple comparisons test.

## Acknowledgements

We gratefully acknowledge support from the Vanderbilt Institute for Nanoscale Science and Engineering (VINSE) for SEM access, the Vanderbilt Technologies for Advanced Genomics (VANTAGE) core for RNA sequencing, and the Vanderbilt Cell Imaging Shared Resource (CISR). This work was funded by the National Science Foundation (NSF) award no. DGE-1937963 (CRD), the National Institutes of Health (NIH) National Institute of Biomedical Imaging and Bioengineering (NIBIB) T32-EB021937 (CRD), R01 EB028690, NIH National Eye Institute (NEI) R01EY022349 (TR, DC, CLD), Marlene & Spencer Hays Directorship (TR), Potoscnak Family Discovery Funds (TR, DC), NIH NEI P30EY008126 (DC), NIH NEI T32EY007135 (AC), and Research to Prevent Blindness Unrestricted Funds (VEI).

## Author Contributions

CRD and SN contributed equally to this work. CRD conducted experiments, acquired and analyzed data, and wrote portions of the manuscript. SN conducted experiments, acquired and analyzed data, and wrote portions of the manuscript. MR acquired and analyzed data and wrote portions of the manuscript. PP conducted experiments and acquired data. TD, AC, SD, RH acquired data. EA and BC conducted experiments. DJC designed study, supervised, provided materials, and edited the manuscript. CLD designed study, supervised, analyzed data, provided materials, and edited the manuscript. TSR designed study, supervised, provided materials, analyzed data, wrote portions of the manuscript and edited the manuscript.

## Data Availability Statement

All data needed to evaluate the conclusions in the paper are present in the paper and/or the Supplementary Materials. RNA sequencing data will be uploaded to GO upon publication.

## MTA

The data can be provided by the corresponding authors pending scientific review and a completed material transfer agreement. Requests for the data should be submitted to.

