## Supplemental Information for "IOP-independent treatment for neurodegeneration in glaucoma using optimized ROS-scavenging microparticles for sustained release of erythropoietin"

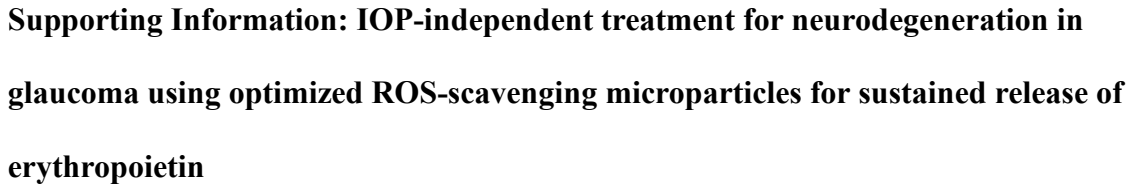

**Supporting Information: IOP-independent treatment for neurodegeneration in
glaucoma using optimized ROS-scavenging microparticles for sustained release of
erythropoietin**

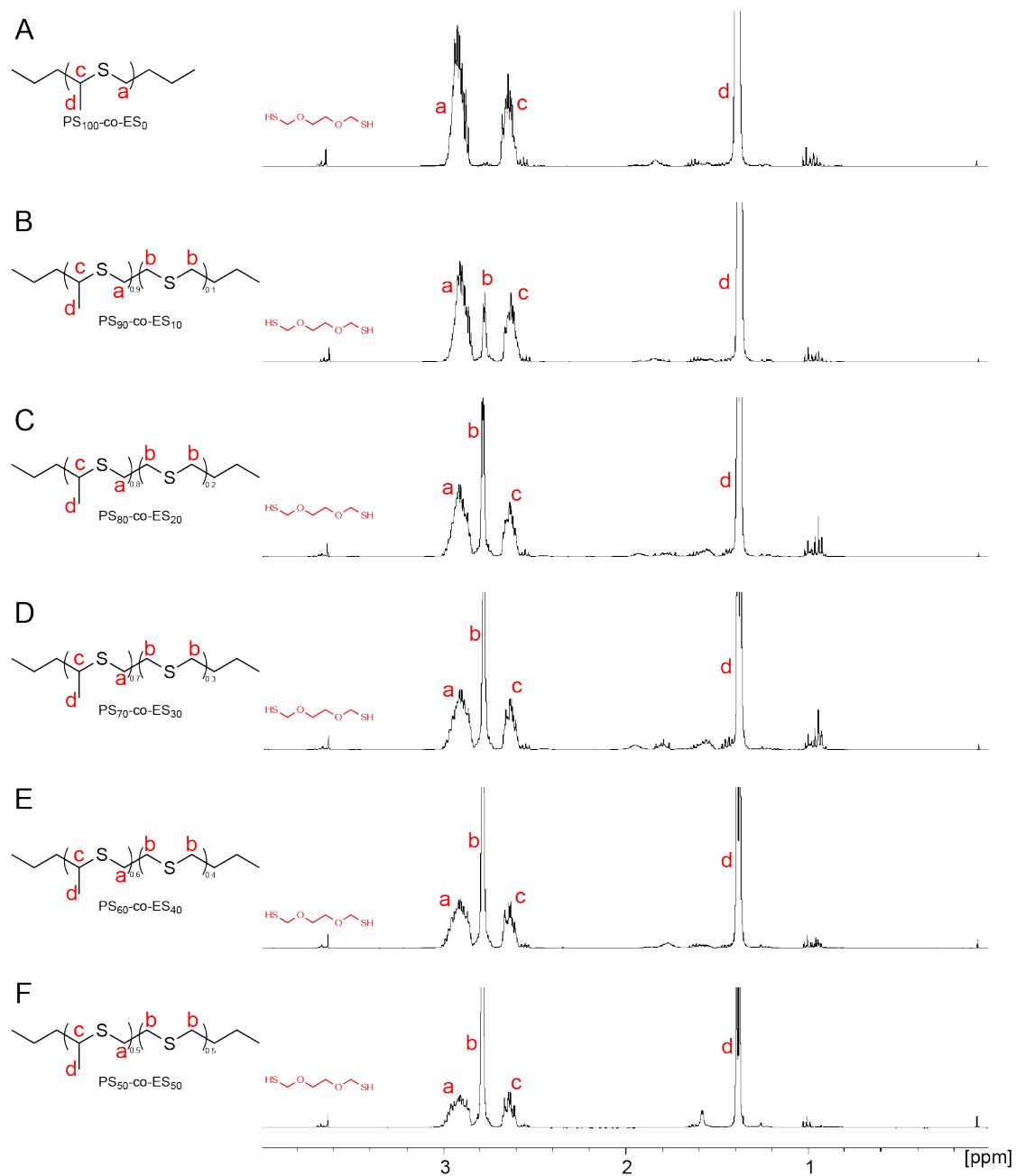

**Supplementary Figure 1:** $^1\text{H}$ Nuclear magnetic resonance (NMR) spectra for the PS-co-ES series. (A) $\text{PS}_{100}\text{-co-ES}_0$; (B) $\text{PS}_{90}\text{-co-ES}_{10}$; (C) $\text{PS}_{80}\text{-co-ES}_{20}$; (D) $\text{PS}_{70}\text{-co-ES}_{30}$; (E) $\text{PS}_{60}\text{-co-ES}_{40}$; (F) $\text{PS}_{50}\text{-co-ES}_{50}$.

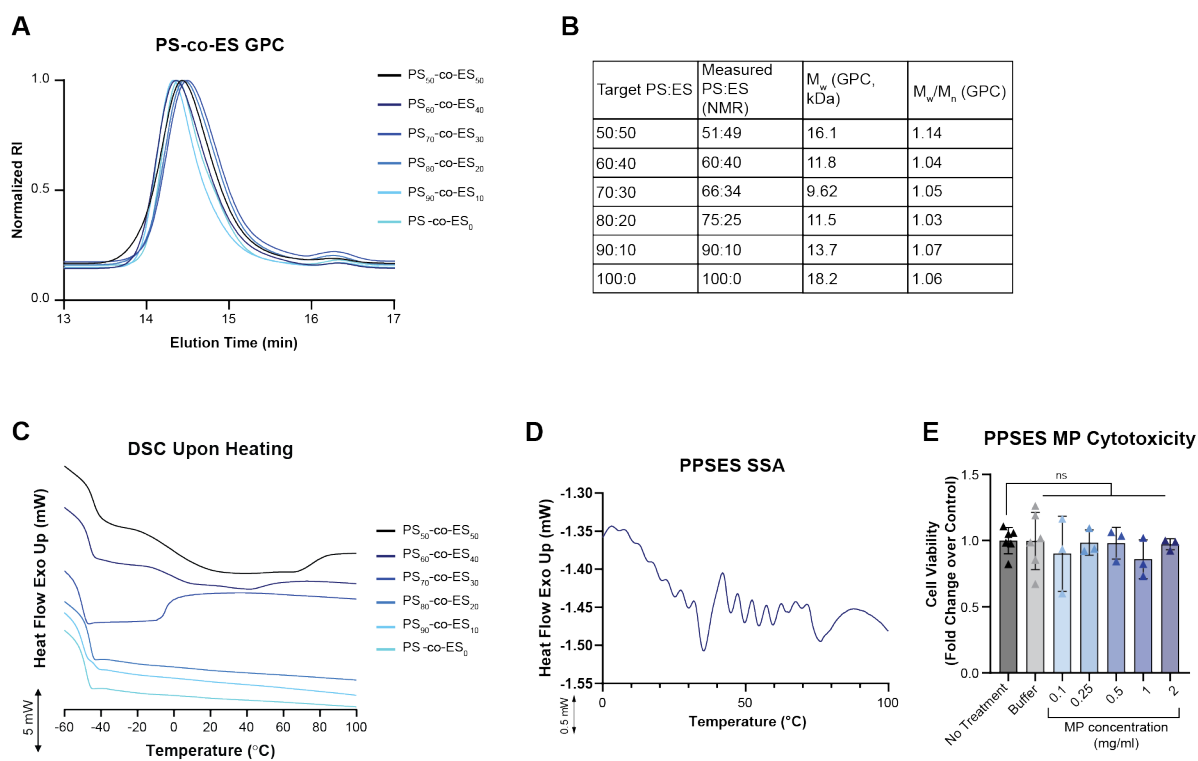

**Supplementary Figure 2:** (A) Gel permeation chromatography (GPC) of copolymer series. (B) Summary table of polymer molecular weight characterization. (C) Differential scanning calorimetry (DSC) as the sample was heated. (D) Successive self-nucleation and annealing (SSA) demonstrating temperature-dependent crystallization of PS<sub>50</sub>-co-ES<sub>50</sub> (PPSES). (E) Viability of ARPE-19 cells after treatment with increasing doses of PPSES microparticles (MP). ES = ethylene sulfide; Exo = exothermic; $M_n$ = number-average molecular weight; $M_w$ = weight-average molecular weight; NMR = nuclear magnetic resonance; PS = propylene sulfide; RI = refractive index.

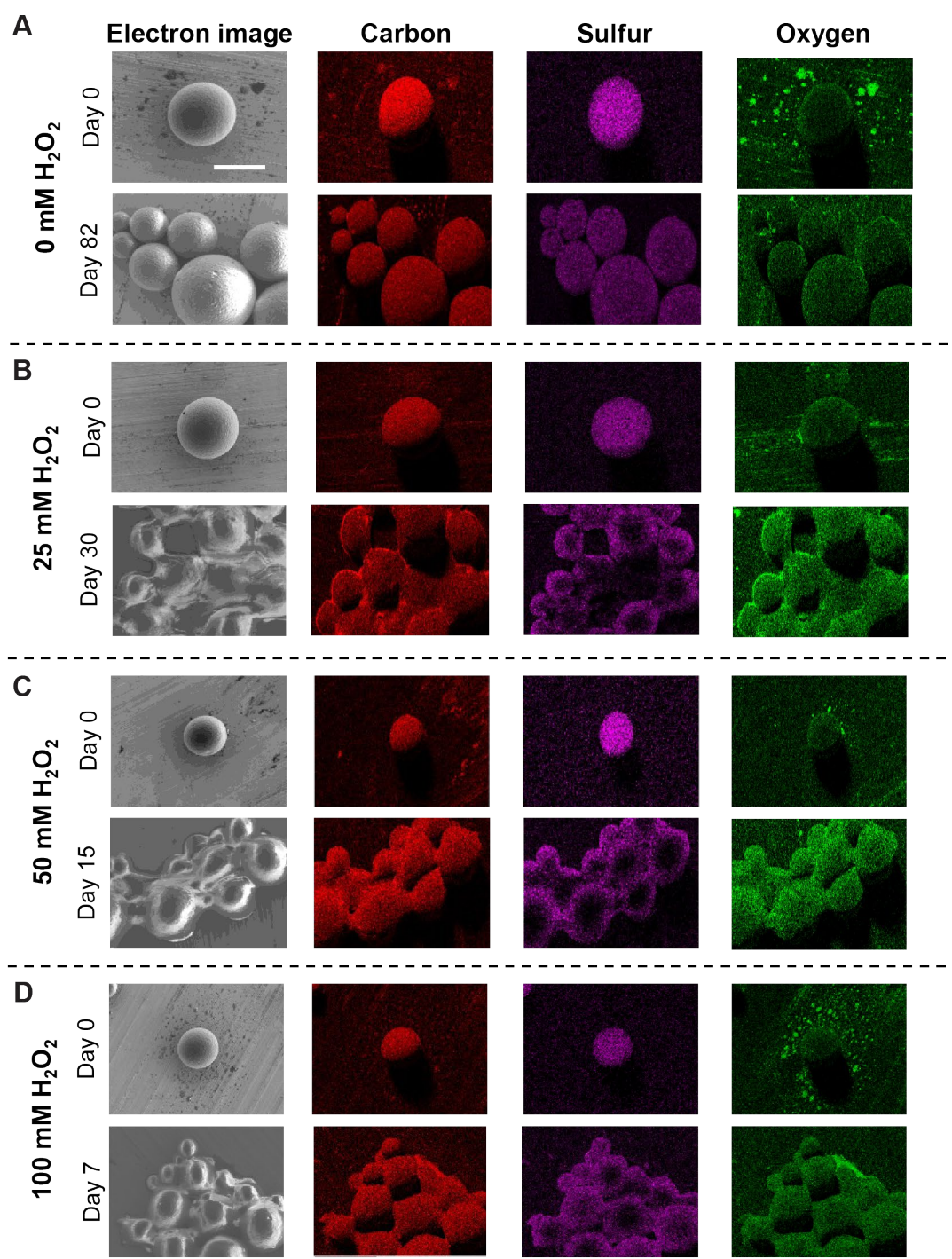

**Supplementary Figure 3:** Elemental mapping of PS<sub>50</sub>-co-ES<sub>50</sub> microparticles (MPs) via x-ray dispersive spectroscopy (EDS) incubated in (A) 0 mM H<sub>2</sub>O<sub>2</sub>, (B) 25 mM H<sub>2</sub>O<sub>2</sub>, (C) 50 mM H<sub>2</sub>O<sub>2</sub>, and (D) 100 mM H<sub>2</sub>O<sub>2</sub> for the indicated amount of time. Scale bar = 100 μm. The time point represents the last day before total degradation and peak oxygen signal at each H<sub>2</sub>O<sub>2</sub> concentration.

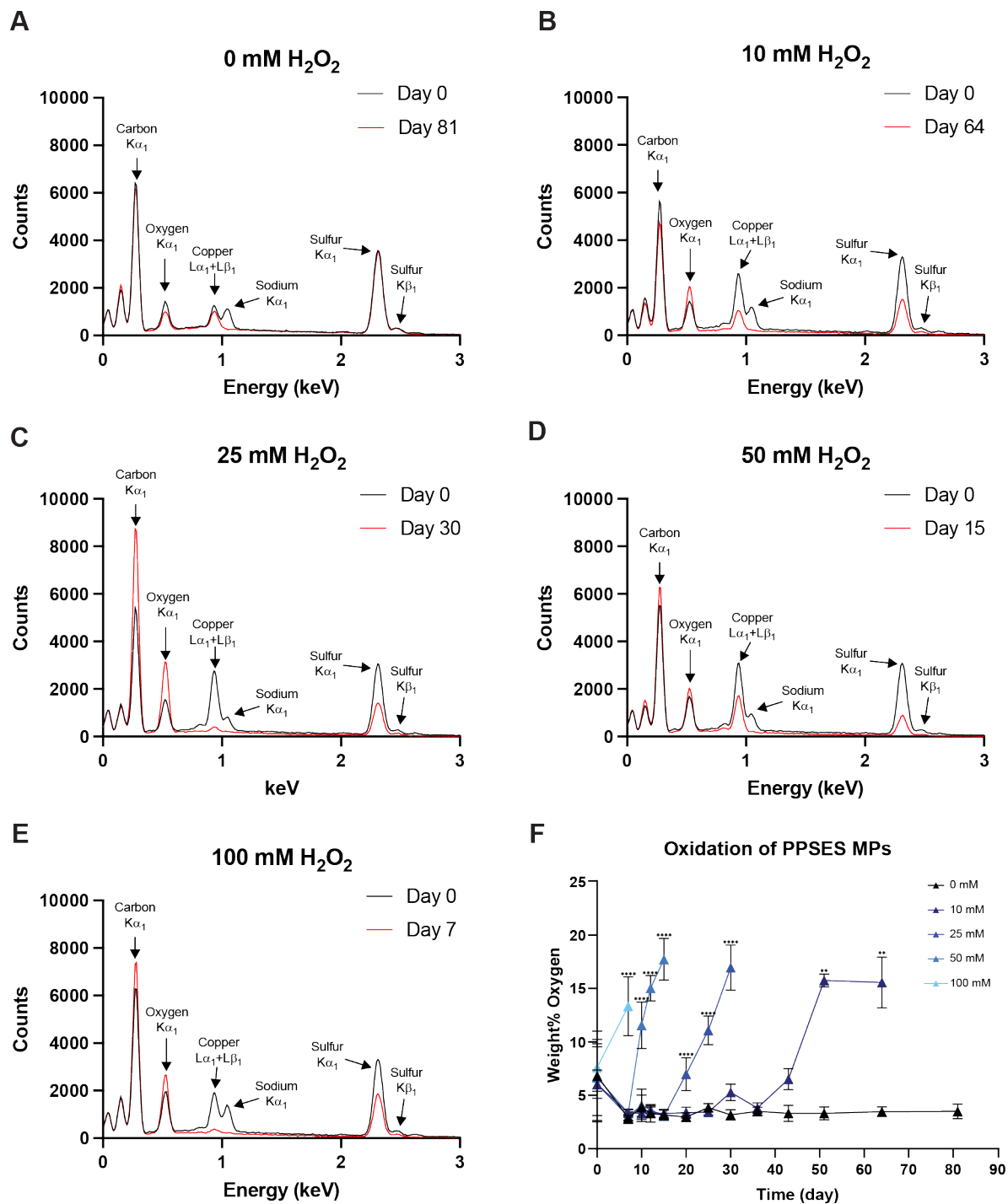

**Supplementary Figure 4:** Representative energy dispersive x-ray spectroscopy (EDS) spectra for PPSES microparticle (MP) oxidation at day 0 and the final time point for each sample. (A) 0 mM $\text{H}_2\text{O}_2$ at day 81; (B) 10 mM $\text{H}_2\text{O}_2$ at day 64; (C) 25 mM $\text{H}_2\text{O}_2$ at day 30; (D) 50 mM $\text{H}_2\text{O}_2$ at day 15; (E) 100 mM $\text{H}_2\text{O}_2$ at day 7. (F) Quantification of oxygen weight % for the time course for each condition (MPs with diameter $<50\ \mu\text{m}$). \*\*$p<0.005$; \*\*\*$p<0.0001$.

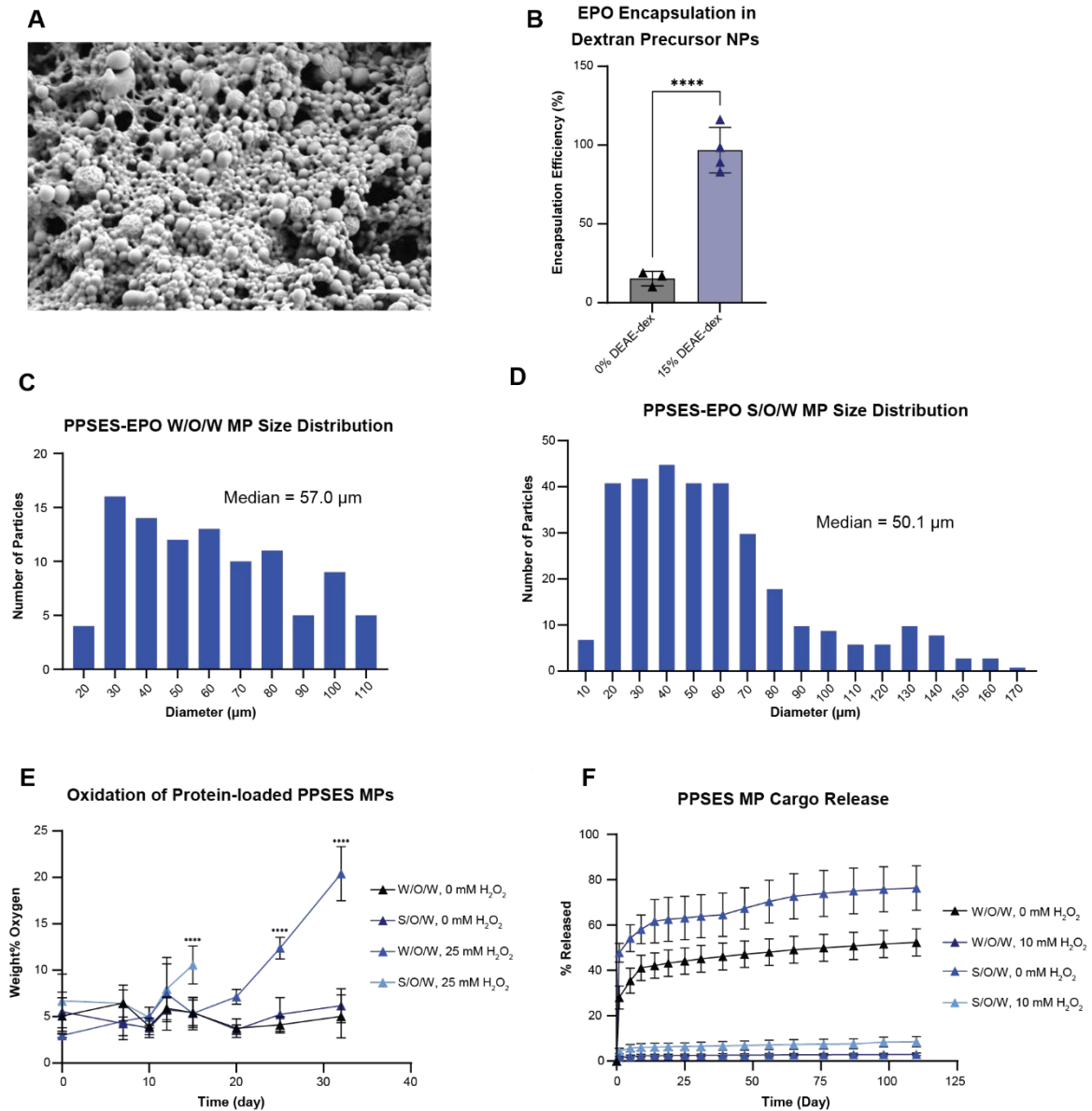

**Supplementary Figure 5:** Characterization of EPO-loaded microparticles (MPs). **(A)** Scanning electron micrograph (SEM) of EPO-loaded diethylaminoethyl (DEAE)-dextran NPs. Scale bar = 2 $\mu\text{m}$. **(B)** Encapsulation efficiency of EPO in DEAE-dextran NPs. **(C-D)** Size distribution of PPSES-EPO MPs formulated by water-in-oil-in-water (W/O/W) emulsion **(C)** and solid-in-oil-in-water (S/O/W) emulsion techniques **(D)**. **(E)** Quantification of oxidation for MPs loaded with the model protein bovine serum albumin (BSA) using the W/O/W and S/O/W techniques. **(F)** Release of rhodamine-tagged BSA from W/O/W and S/O/W formulations. \*\*\*\*$p < 0.0001$.

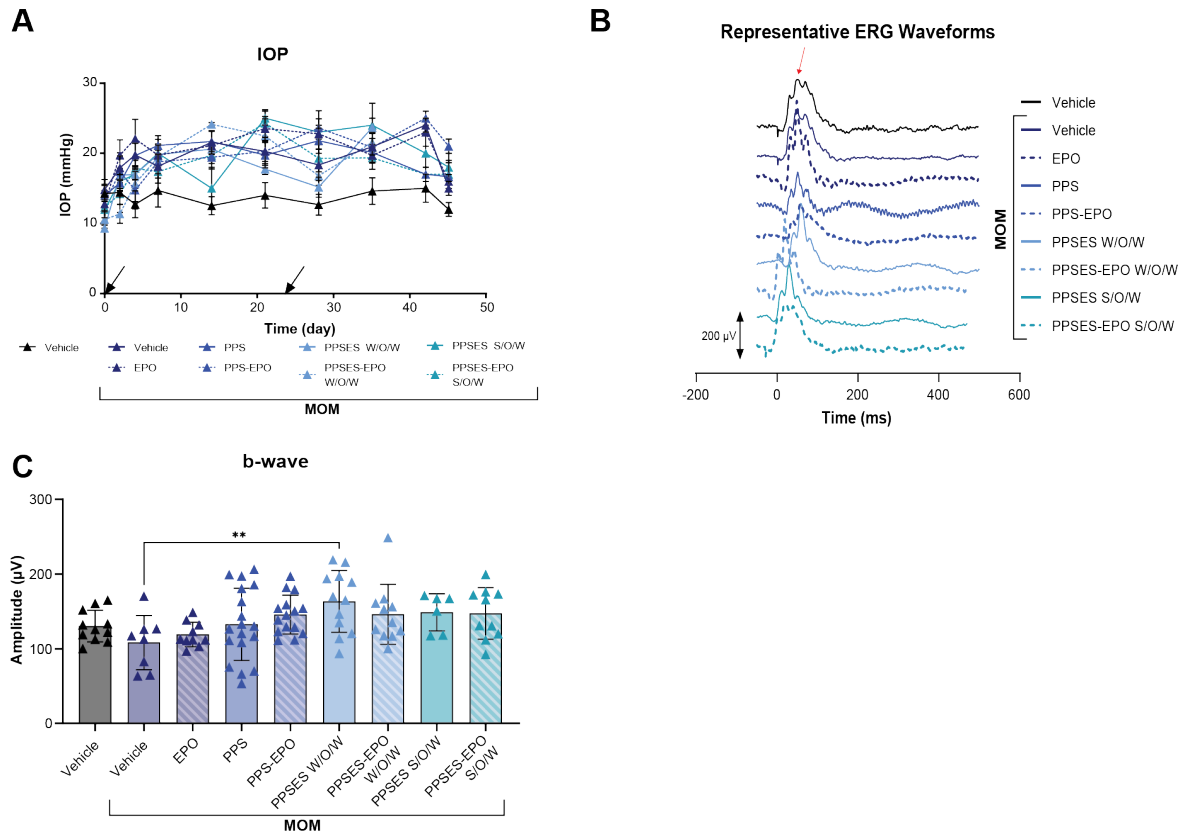

**Supplementary Figure 6:** (A) Intraocular pressure (IOP) measurements over the course of the study. Arrows indicate the day of microbead injections. (B) Representative electroretinogram (ERG) waveforms with arrow indicating amplitude measurement. (C) Quantification of ERG b-wave amplitude. \*\*$p < 0.005$. MOM=microbead occlusion model.

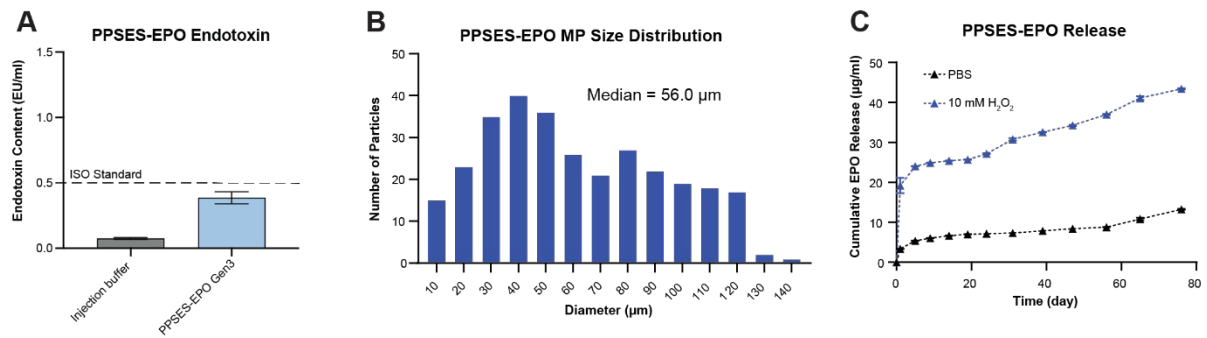

**Supplementary Figure 8:** Characterization of PPSES-EPO microparticles (MPs) optimized for non-human primate application. **(A)** Endotoxin levels in PPSES-EPO MPs. **(B)** Size distribution for PPSES-EPO MPs. **(C)** *In vitro* release of rhodamine-labeled EPO.

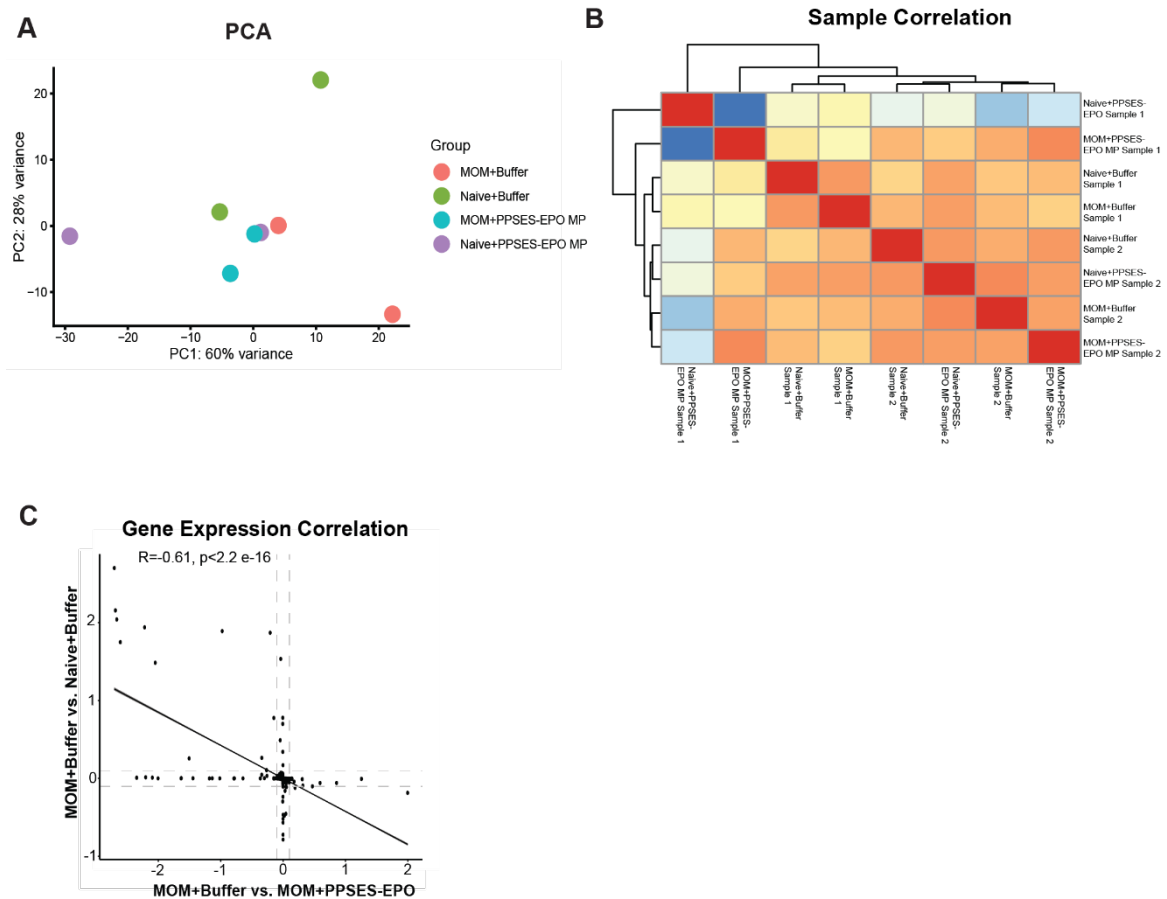

**Supplementary Figure 8:** **(A)** Principal component analysis (PCA). Each dot represents one bulk RNA-seq sample generated from 2–3 pooled retinas to obtain sufficient RNA for sequencing. **(B)** Heatmap showing correlations between each biological sample. **(C)** Correlation between genes in MOM+Buffer vs. Naive+Buffer and MOM+Buffer vs. MOM+PPSES-EPO MP. MOM = microbead occlusion model.

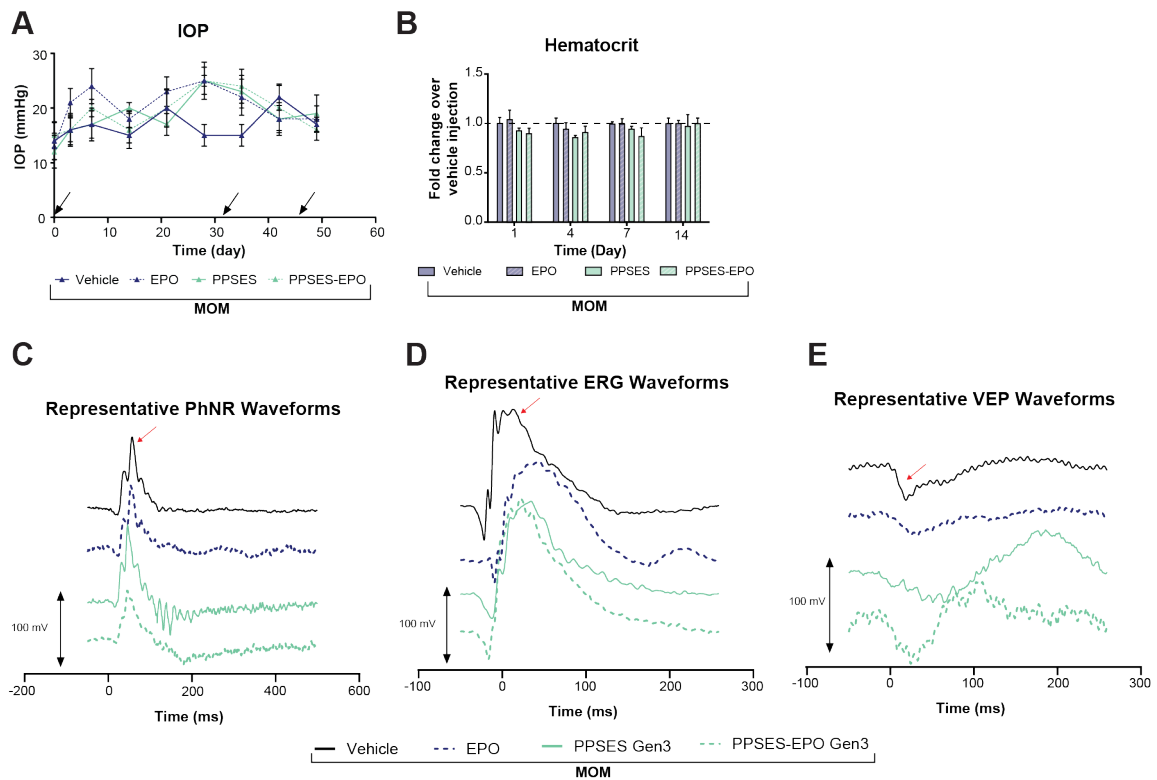

**Supplementary Figure 9:** (A) Intraocular pressure (IOP) throughout the study. Arrows indicate days when microbeads were injected. (B) Hematocrit measures normalized to buffer-injected animals (dotted line). (C-E) Representative waveforms for (C) photopic negative response (PhNR), (D) electroretinogram (ERG), and (E) visual evoked potential (VEP) 7 weeks after first microbead injection.

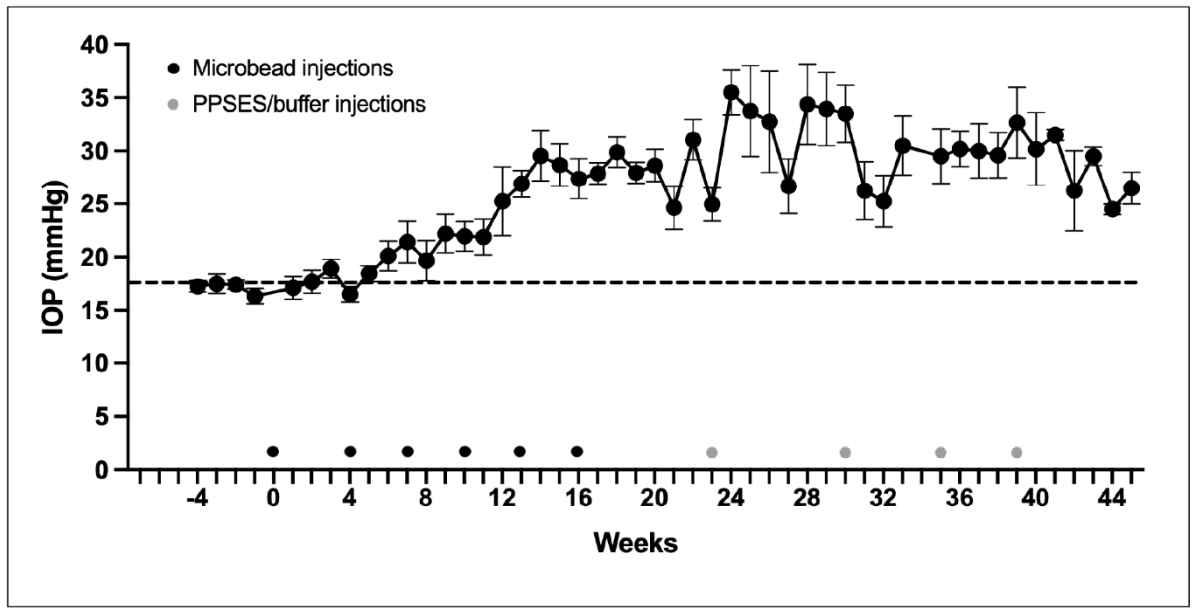

**Supplementary Figure 10:** Intraocular pressure (IOP) measurements in awake non-human primates (NHPs) before and after intracameral microbead injection (black dots). Grey dots indicate averaged times of intravitreal injections of buffer, PPSES, or PPSES-EPO microparticles (MPs). Error bars represent SEM. Dotted line represents the pre-microbead, normative IOP level for the NHPs.

**Supplementary Table 1:** Non-human primate IFN-γ concentration (aqueous humor)

| Treatment | Week | Eye | IFN-γ (pg/ml) |
| --- | --- | --- | --- |
| PPSES-EPO | 0 | RE | 4.8 |
|  |  | LE | 4.0 |
|  | 6 | RE | 3.8 |
|  |  | LE | 4.5 |
|  | 10 | RE | 4.9 |
|  |  | LE | 3.8 |
|  | 15 | RE | 4.8 |
|  |  | LE | 4.0 |
|  | 20 | RE | 4.8 |
|  |  | LE | 3.5 |
|  | 0 | RE | 4.3 |
|  |  | LE | 4.3 |
|  | 4 | RE | 4.0 |
|  |  | LE | 4.9 |
| PPSES-blank | 9 | RE | 4.0 |
|  |  | LE | * |
|  | 11 | RE | 3.8 |
|  |  | LE | * |
|  | 0 | RE | 3.5 |
|  |  | LE | 3.5 |
|  | 6 | RE | 5.0 |
|  |  | LE | 4.2 |
|  | 10 | RE | 4.5 |
|  |  | LE | 3.5 |
|  | 15 | RE | 3.0 |
|  |  | LE | 4.0 |
|  | 20 | RE | 2.5 |
|  |  | LE | 3.5 |

\*Eye exhibited abnormally high intraocular pressure

**Supplementary Table 2:** Murine study #1 (Endpoint 6-7 weeks after initial MOM)
Figures 2-3 and Supplementary Figure 6

| Treatment Group | N (# of eyes) | EPO Dose (ng) |
| --- | --- | --- |
| Naive + Vehicle | 12 | N/A |
| MOM + Vehicle | 14 | N/A |
| MOM + EPO | 12 | 8.5 |
| MOM + PPS | 16 | N/A |
| MOM + PPS-EPO | 18 | 8.5 |
| MOM + PPSES W/O/W | 16 | N/A |
| MOM + PPSES-EPO W/O/W | 14 | 7.7 |
| MOM + PPSES S/O/W | 8 | N/A |
| MOM + PPSES-EPO S/O/W | 12 | 7.3 |

**Supplementary Table 3:** Murine study #2 (Endpoint 3 weeks after initial MOM)
Supplementary Figure 7

| Treatment Group | N (# of eyes) | EPO Dose (ng) |
| --- | --- | --- |
| Naive + Vehicle | 5 | N/A |
| MOM + Vehicle | 5 | N/A |
| Naive + PPSES-EPO | 5 | 102 |
| MOM + PPSES-EPO | 5 | 102 |

**Supplementary Table 4:** Murine study #3 (Endpoint 9 weeks after initial MOM)
Figure 4 and Supplementary Figure 9

| Treatment Group | N (# of eyes) | EPO Dose (ng) |
| --- | --- | --- |
| MOM + Vehicle | 12 | N/A |
| MOM + EPO | 16 | 100 |
| MOM + PPSES | 20 | N/A |
| MOM + PPSES-EPO | 20 | 102 |

**Supplementary Table 5:** Summary of non-human primate treatment groups

| Treatment Group | N (# of eyes) | EPO Dose (ng) |
| --- | --- | --- |
| MOM + Vehicle | 3 | N/A |
| MOM + PPSES | 4 | N/A |
| MOM + PPSES-EPO | 4 | 1000 |
| MOM + PPSES-EPO<br>(rhodamine-tagged) | 1 | 1000 |
